# A multivariate method to correct for batch effects in microbiome data

**DOI:** 10.1101/2020.10.27.358283

**Authors:** Yiwen Wang, Kim-Anh Lê Cao

## Abstract

Microbial communities are highly dynamic and sensitive to changes in the environment. Thus, microbiome data are highly susceptible to batch effects, defined as sources of unwanted variation that are not related to, and obscure any factors of interest. Existing batch correction methods have been primarily developed for gene expression data. As such, they do not consider the inherent characteristics of microbiome data, including zero inflation, overdispersion and correlation between variables. We introduce a new multivariate and non-parametric batch correction method based on Partial Least Squares Discriminant Analysis. PLSDA-batch first estimates treatment and batch variation with latent components to then subtract batch variation from the data. The resulting batch effect corrected data can then be input in any downstream statistical analysis. Two variants are also proposed to handle unbalanced batch x treatment designs and to include variable selection during component estimation. We compare our approaches with existing batch correction methods removeBatchEffect and ComBat on simulated and three case studies. We show that our three methods lead to competitive performance in removing batch variation while preserving treatment variation, and especially when batch effects have high variability. Reproducible code and vignettes are available on GitHub.

## Introduction

Investigating the link between microbial composition and phenotypes, including human diseases has become critical in microbiome research. The microbiome was first defined as the microorganisms and their activities within their specific habitats (Prescott, 2017) then widely referred to as the genetic material within the entire collection of microorganisms. The microbiome can be considered as a counterpart to the human genome, as microbes include a large population and participate in human physiological system, such as programming the immune system and providing nutrients. The disruption of gut microbial communities has been linked to varieties of diseases and sub-health status, ranging from inflammatory bowel diseases (Zuo and Ng, 2018), diabetes (Sharma and Tripathi, 2019) to obesity (Gérard, 2016) and malnutrition (Tidjani Alou *et al*., 2017).

Microbiome research faces the challenges of data reproducibility and replicability that are essential to the validity of the statistical results. In particular, microbial communities are highly dynamic (Schloss, 2018) and thus microbiome data are highly susceptible to batch effects, that is, any unwanted sources of variation that are unrelated to and obscure the biological factors of interest (Wang and Lê Cao, 2019). Microbiome studies affected by batch affects are abundant in the literature: For example, unwanted variation can be introduced by sequencing batches (Hieken *et al*., 2016) or independent studies (Duvallet *et al*., 2017). Other confounding factors including geography, age, sex, health status, stress and diet also introduce batch effects to the composition of the host microbiota (Gibson *et al*., 2004, Lozupone *et al*., 2013, Haro *et al*., 2016, Kim *et al*., 2017).

Two types of approaches exist to handle batch effects (Wang and Lê Cao, 2019): methods that correct for batch effects consist in removing batch variation from the data, while methods that account for batch effects include batch effects as covariates in the statistical model. Correcting for batch effect offers the flexibility to apply any type of downstream analysis, including dimension reduction, visualisation and clustering. Methods accounting for batch effects are often restricted to differential abundance analysis with models that hold strong assumptions about data distribution, they include zero-inflated Gaussian model (Paulson *et al*., 2013) and Bayesian Dirichlet multinomial regression (Dai *et al*., 2018)). In terms of evaluating the effectiveness of batch effect handling methods, batch effect removal methods are more straightforward and transparent than methods that account for batch effects. However, correcting for batch effects in microbial studies is challenging. Microbiome studies usually include a small sample size, which increases the uncertainty of batch effect estimation (Debelius *et al*., 2016). The data also have inherent characteristics including zero inflation, uneven library sizes, compositional structure and inter-variable dependency which challenge existing batch effect correction methods such as ComBat (Johnson *et al*., 2007) and removeBatchEffect (Ritchie *et al*., 2015) that were developed for gene expression data. While methods accounting for batch effects consider microbiome data characteristics within models, batch effect correction methods are often applied to microbiome data that are transformed to meet the methods’ parametric assumptions. However, such transformations are not sufficient to address zero-inflated distribution and the variables’ inter-dependency.

As microbiome data are naturally multivariate, univariate methods such as removeBatchEffect and ComBat are limited and do not take into account the microbial variables inter-dependency (Ramette, 2007). Another limitation of existing methods (e.g., ComBat) is their assumption that batch effects are systematic and thus have a homogeneous influence on all variables. However, batch effects in microbiome data were found to be non-systematic (Wang and Lê Cao, 2019). When non-systematic batch effects are mistakenly treated as systematic, biological variation of interest might be removed, or the batch variation may remain during the batch effect correction process. The multivariate method Remove Unwanted Variation (RUV) has been recently adapted for microbiome data (Hardwick *et al*., 2018, Moskovicz *et al*., 2020), but requires the availability of negative control variables and technical sample replicates that capture batch variation, which are not often available in microbiome studies.

Finally, another challenge that batch effect correction methods face is their assumption that batch and treatment effects are independent, requiring a balanced batch × treatment design (Wang and Lê Cao, 2019). However, technical experimental issues result in unbalanced designs, where batch and treatment effects are confounded, resulting in losing treatment variation during the batch correction process. Promising methods have been proposed in other fields of application, such as single cell RNA-seq field. Methods such as Seurat V3 (Stuart *et al*., 2019), mnnCorrect (Haghverdi *et al*., 2018), scmerge (Lin *et al*., 2019), zinbwave (Risso *et al*., 2018) assume a zero-inflated distribution but are not directly applicable to microbiome studies because of their very small sample size compared to the single cell datasets.

We propose a novel method to correct for batch effects in microbiome data based on Partial Least Squares Discriminant Analysis (PLSDA, Barker and Rayens 2003). Our approach, PLSDA-batch is highly suitable for microbiome studies as it is non-parametric, multivariate and allows for ordination. It estimates latent components related to treatment and batch effects to remove batch variation in the data whilst preserving biological variation of interest. Two variants are proposed to 1/ handle unbalanced batch × treatment designs and 2/ select discriminative microbial variables amongst treatment groups. We assess the performance of PLSDA-batch in simulation and three case studies which investigate microbial communities in sponge tissues, anaerobic digestion conditions and diet types. We compare the efficiency of our approaches in removing batch effects and uncovering treatment effects with ComBat and removeBatchEffect.

## Methods

PLSDA-batch is derived from Partial Least Squares Discriminant Analysis (Barker and Rayens, 2003). We first give a brief description of PLSDA and its core method Partial Least Squares (PLS, Wold *et al*. 2001). We will use the following notations: ***X*** denotes an (*n* × *p*) explanatory data matrix with *p* microbial variables and ***Y*** an (*n* × *q*) data matrix with *q* response variables. Both datasets match on the same n samples. We denote the matrix transpose by ^T^. The *ℓ*_1_ norm of a random vector 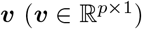 is defined as 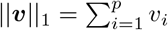 and the *ℓ*_2_ norm is 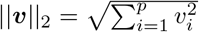.

### Partial Least Squares

PLS, a.k.a Projection to Latent Structures is an orthogonal component-based regression method commonly used to model the covariance structure between explanatory (***X***) and response (***Y***) matrices in large datasets. The optimisation problem to solve is:

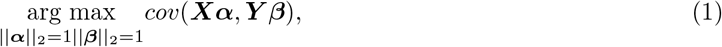

where 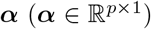 and 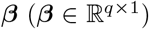 represent the loading vectors of ***X*** and ***Y*** respectively. The aim of PLS is to find the linear transformations (***α*** and ***β***) of ***X*** and ***Y*** that maximise the covariance between their latent components denoted as ***t*** and ***u*** respectively, with ***t*** = ***Xα*** and 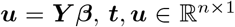. After the first pair of latent components (***t, u***) is obtained, the residual matrix is calculated via matrix deflation:

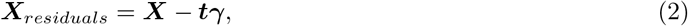

where ***γ*** = (***t***^T^***t***)^-1^***t***^T^***X***. ***γ*** represents the regression coefficient vector for each variable in ***X*** on ***t***, 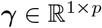. Similarly, we can calculate the residual matrix ***Y**_residuals_* by deflating the matrix ***Y*** with ***u***. The deflated matrices are then used as updated ***X*** and ***Y*** for the next PLS dimension. The deflation steps ensure that the latent components associated to each PLS dimension are orthogonal.

### PLS Discriminant Analysis

PLSDA is an adaption of PLS for classification and discrimination, where the response matrix ***Y*** is a dummy matrix transformed from a categorical outcome variable. Each column in ***Y*** indicates the group membership of samples: If sample i belongs to group *j*, then *Y_ij_* is 1, otherwise 0. For each dimension *h* =1,…, *H*, the latent components ***t**_h_* and ***u**_h_* are calculated as shown earlier in Eq.(1) of the section “Partial Least Squares”. ***t**_h_* summarises the variation from ***X*** that is associated with ***u**_h_*, where ***u**_h_* is a linear combination of the dummy outcomes in ***Y***. Thus, the ***t**_h_* component is mostly relevant to explain the discrimination between sample groups.

In PLSDA, we need to specify the optimal number of components *H*. It can be chosen using repeated cross-validation to estimate the classification error rate on each component ***t**_h_*. As PLSDA is an iterative process based on deflated matrices, the *H* components that yield the lowest error rate correspond to the overall performance of the PLSDA model (Rohart *et al.*, 2017).

### sparse PLSDA

sPLSDA uses *ℓ*_1_ penalisation on the loading vectors [***α***_1_,…, ***α**_H_*] in PLSDA to select variables (Lê Cao *et al*., 2011). During the regression step, for each component *h* = 1,…, *H*, the penalty is solved with soft-thresholding in Eq.(1):

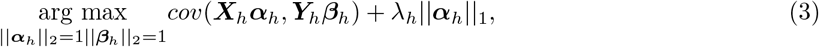

where *λ_h_* is a non-negative parameter that controls the amount of shrinkage on the loading vector ***α**_h_* and thus the number of non-zero loadings. The latent component ***t**_h_* is therefore calculated based on a subset of variables that are deemed the most discriminative to classify the sample groups.

Two types of parameters need to be specified in sPLSDA: the number of components *H* and the number of variables to select on each component, which corresponds to the shrinkage coefficient *λ_h_*. Both parameters can be chosen simultaneously using repeated cross-validation by evaluating the classification error rate on a grid of number of variables to select on each component (Rohart *et al*., 2017).

### PLSDA-batch

PLSDA-batch aims to estimate and remove batch variation whilst preserving treatment variation. We use additional notations as we include in the model two different types of sample information, treatment and batch, denoted ***Y***^(*trt*)^ and ***Y***^(*batch*)^ respectively. The matrices 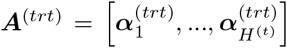 and 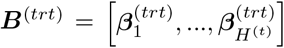 include the loading vectors associated to ***X*** and ***Y***^(*trt*)^ respectively, where *H*^(*t*)^ is the number of components associated to the treatment variation. The corresponding latent components are denoted 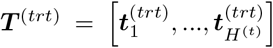 and 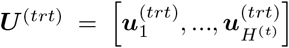. Similar notations are used for the loading vectors and latent components associated to the batch effect across *H*^(*b*)^ components. We will use simplified notations without superscript, such as ***Y***, ***A***, *H* and ***T*** that are related to either treatment or batch variation when there is not ambiguity. ***X***^(*nobatch*)^ is the matrix from which the batch effect is removed, and similarly ***X***^(*notrt*)^ for the treatment effect.

#### Overview

The general concept of PLSDA-batch is shown in the first column of Figure 1. Assuming ***X*** includes both treatment and batch effects, the samples projected onto a Principal Component Analysis (PCA) plot would be segregated according to both treatment and batch information. In a first step, PLSDA-batch estimates the treatment variation via the components ***T***^(*trt*)^, which are extracted out of ***X*** to obtain ***X***^(*notrt*)^. Thereafter, only the batch variation still remains. The second step estimates the batch associated components ***T***^(*batch*)^ from ***X***^(*notrt*)^. The original dataset ***X*** is then deflated with ***T***^(*batch*)^ to obtain the final matrix corrected for batch effects whilst preserving the treatment variation ***X***^(*nobatch*)^.

**Figure 1:**
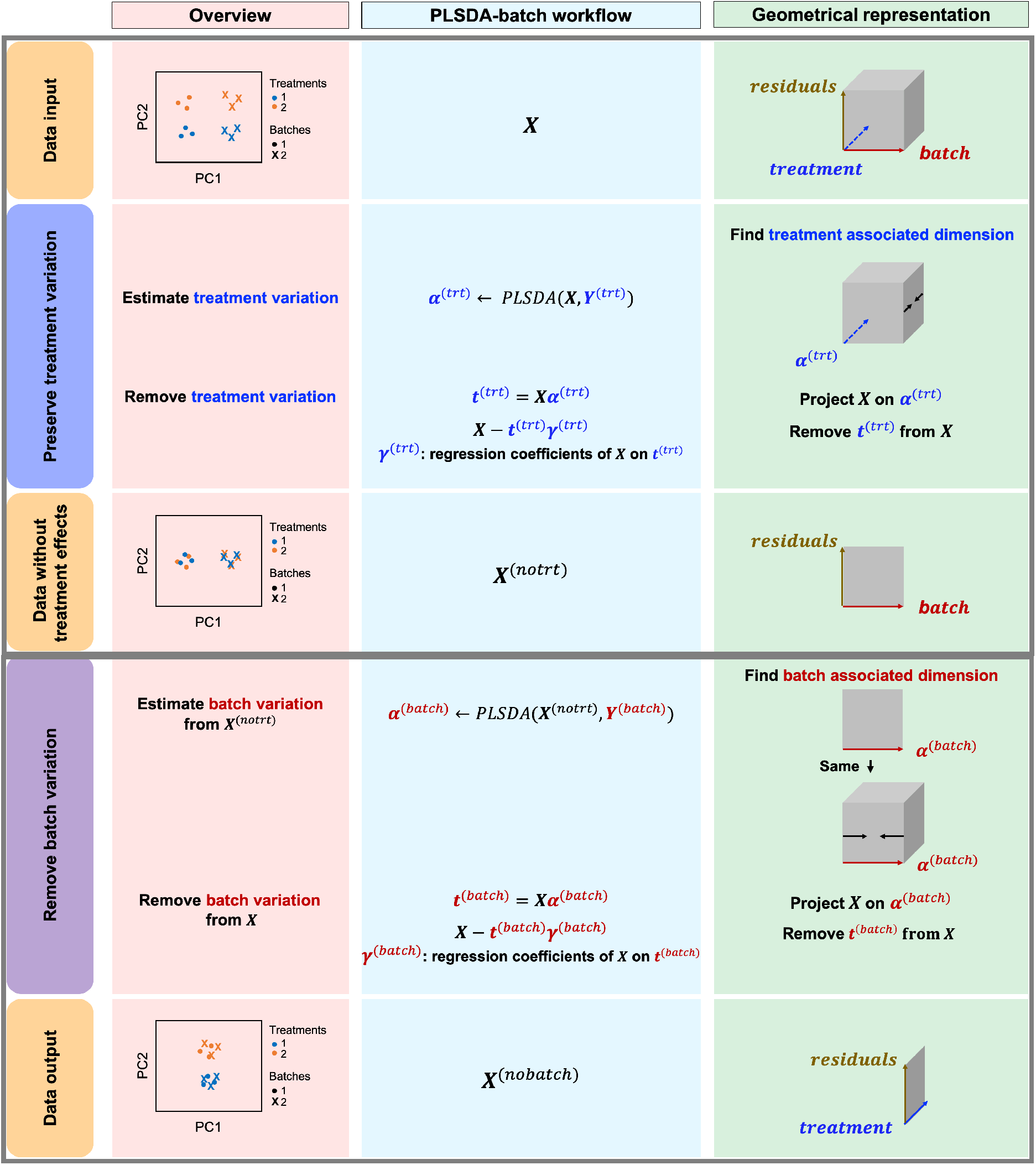
PLSDA-batch framework. From left to right columns: Visualisation with Principal Component Analysis sample plots; Workflow describing each step of Algorithm 1 and Geometrical representation of the approach via projections and deflation. For illustrative purpose, we only represent one component associated with either treatment or batch effects.

#### Algorithmic and geometrical point of views

The remaining columns in Figure 1 further describe the approach. For illustrative purposes, we only depict the case where one component is associated with either treatment or batch effects rather than several components. The data matrix ***X*** with both treatment and batch effects can be decomposed into three major sources of variation: treatment, batch and residuals, which are assumed to be orthogonal. In practice however, treatment and batch sources are likely to be correlated to some extent, which motivated our approach to first estimate the treatment variation to avoid over-estimating the batch variation and losing substantial treatment variation.

In the first step, we apply PLSDA to ***X*** and ***Y***^(*trt*)^ to identify the dimension of treatment effects ***α***^(*trt*)^ from ***X*** (see Algorithm 1 “Estimation of latent dimensions”). ***t***^(*trt*)^ is then calculated using a scalar projection of ***X*** onto ***α***^(*trt*)^. Therefore, the treatment variation of all variables in ***X*** is summarised in the component ***t***^(*trt*)^. We then calculate the matrix without treatment effects ***X***^(*notrt*)^ by deflating ***X*** with **t**^(*trt*)^. In the second step, we identify the batch associated dimension ***α***^(*batch*)^ from ***X***^(*notrt*)^, then calculate ***t***^(*batch*)^ by projecting ***X*** onto ***α***^(*batch*)^. The batch variation *t*^(*batch*)^ is then removed from ***X*** via *matrix deflation* whilst ensuring the treatment effects are fully preserved. Since the components ***t***^(*trt*)^ and ***t***^(*batch*)^ are orthogonal, we could also deflate ***X***^(*notrt*)^ with respect to ***t***^(*batch*)^ but such alternative would require adding the treatment variation back.

#### Weighted PLSDA-batch

A balanced batch × treatment design is an experimental design where samples within each treatment group are evenly distributed across batches (Wang and Lê Cao, 2019). Because of quality control steps or lack of samples, a batch × treatment design may be unbalanced, resulting in treatment and batch effects that are correlated and not separable. In our approach, latent components associated to either treatment or batch effects are orthogonal, which limit our ability to consider the correlation between these two effects. The consequences might be over-estimation of the treatment variation as well as insufficient removal of the batch variation. Weighted PLSDA-batch (wPLSDA-batch) is inspired from weighted PCA to account for unbalanced designs (Holmes and Huber, 2018). We weight each sample *i* with *w_i_* to take into account the number of samples within each batch and treatment:

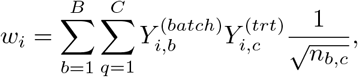

where 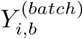 represents the indicator value (0 or 1) of sample i and batch *b* in the dummy matrix ***Y***^(*batch*)^, and similarly for 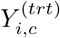. *n_b,c_* represents the sample size in batch b and treatment group *c*. ***W*** is a diagonal matrix that includes *w_i_, i* = 1,…, *n*. We then obtain the weighted explanatory and response matrices ***X***^(*weighted*)^ and ***Y***^(*weighted*)^ multiplying ***X*** and ***Y*** by ***W*** respectively. The batch effect corrected data ***X***^(*nobatch* & *weighted*)^ resulting from the calculation on the weighted matrices using PLSDA-batch are then multiplied by ***W***^-1^ to remove the influence of weights.

#### sparse PLSDA-batch

In PLSDA-batch, the latent components are calculated based on all variables. However, we should assume that treatment effects only affect a small number of variables, while batch effects that include a high variability should affect a large number of variables. A non-sparse version of PLSDA-batch may hence result in treatment associated components ***T***^(*trt*)^ that include the variation from batch related variables, and ultimately affect the accuracy of the batch corrected matrix ***X***^(*nobatch*)^.

To avoid overfitting when we estimate the treatment associated components, we apply *ℓ*_1_-penalty to each loading vector (see Eq. (3)) to select variables. Thus, the variables with no treatment effect are assigned a zero loading value and are not included in the calculation of a component. As we assume that batch effects are more variable than treatment effects, variable selection is not considered when estimating the batch components to ensure that all batch variation is retained.

**Algorithm 1.**
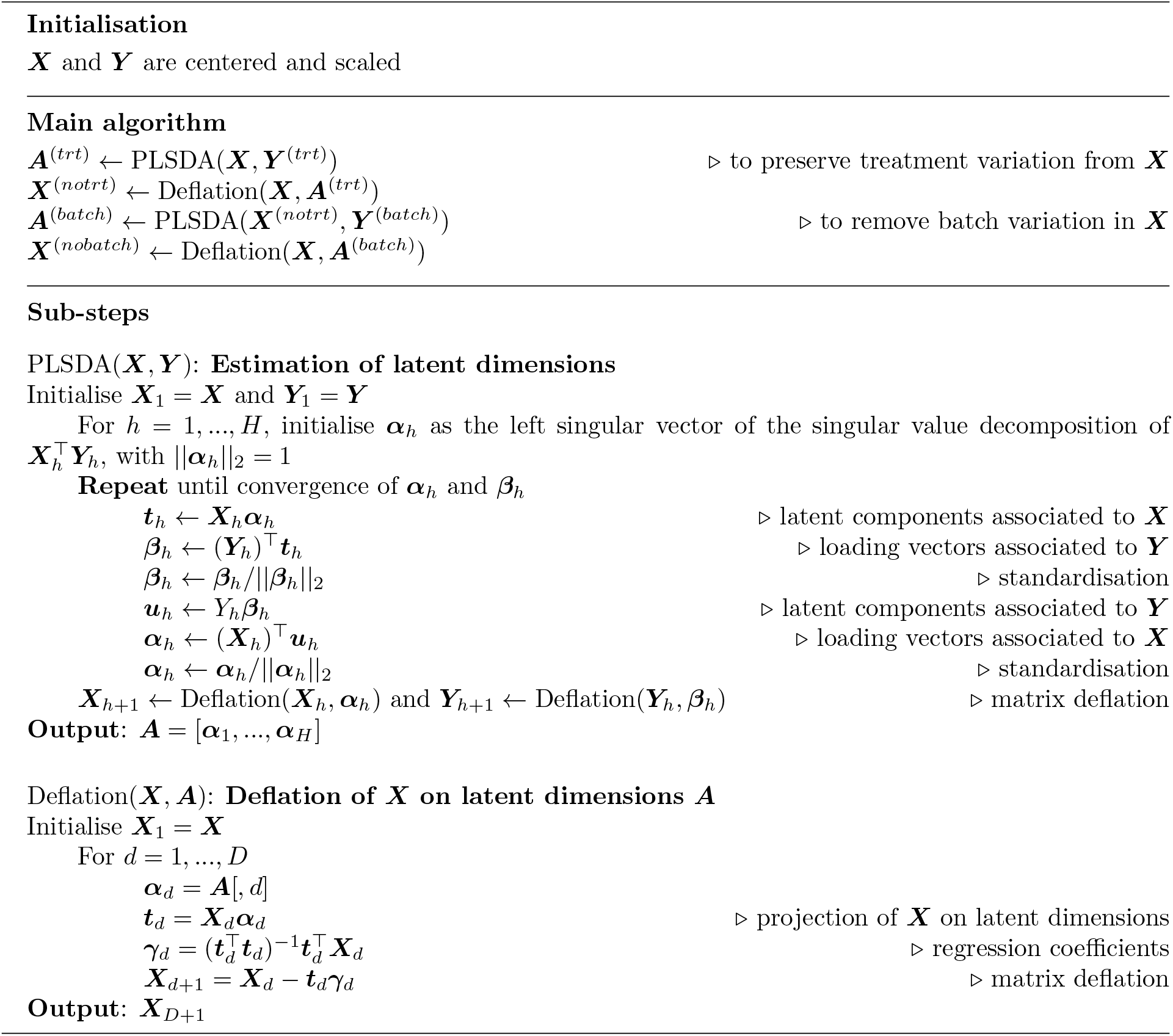
PLSDA-batch.

#### Parameter tuning

In PLSDA-batch, we need to specify the optimal number of components associated with either treatment or batch effects (*H*^(*t*)^ or *H*^(*b*)^). To choose this parameter, we estimate the variance explained in the outcome matrix ***Y***^(*trt*)^ on each treatment component 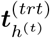, *h*^(*t*)^ = 1,…, *H*^(*t*)^ and similarly for the batch associated outcome matrix and components. We choose the optimal number of components that explain 100% variance in ***Y***, either ***Y***^(*trt*)^ or ***Y***^(*batch*)^. The remainder components should only explain some (unknown) noise.

In sPLSDA-batch, in addition to the number of components, we also need to specify the optimal number of variables to select on each treatment component. For this purpose, we calculate the balanced classification error rate 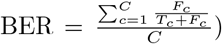, where *F_c_* and *T_c_* represent the number of false and truly classified samples in the treatment group *c, c* = 1,…, *C*, where *C* represents the total number of treatment groups (Tharwat, 2018). The BER is evaluated through repeated cross-validation using the “maximum” prediction distance as described in Rohart *et al*. (2017) on a proposed grid of numbers of variables to select on each treatment component. The number of variables with the lowest BER has the strongest association with the treatment information (***Y***^(*trt*)^).

### Simulation and case studies

#### Simulation study

We adapted the simulation strategy that is component-based and multivariate from Singh *et al*. (2019). We assumed the input data are Centered Log Ratio (CLR) transformed with a Gaussian-like distribution (see section “Case studies”). Thus, we simulated components from a Gaussian distribution across all samples. The data matrix was generated based on the simulated components and corresponding loading vectors for each variable. Different parameters including amount of batch and treatment variability, number of variables with batch and/or treatment effects, balanced and unbalanced batch × treatment designs were considered and summarised in Table 2.

**Table 1:** **Unbalanced batch × treatment design** in the simulation study

|  | Trt1 | Trt2 |
| --- | --- | --- |
| Batch1 | 4 | 16 |
| Batch2 | 16 | 4 |

**Table 2:** Summary of simulations. For a given choice of parameters listed, each simulation was repeated 50 times. *p*^(*trt*)^, *p*^(*batch*)^ and *p*^(*trt* & *batch*)^ represent the number of variables with treatment, batch, or both effects respectively. **Simulation 6** includes parameters reflective of real data.

| Parameters | $\mu_{(trt)}$ | $\sigma_{(trt)}$ | $\mu_{(batch)}$ | $\sigma_{(batch)}$ | $p^{(trt)}$ | $p^{(batch)}$ | $p^{(trt \& batch)}$ |
| --- | --- | --- | --- | --- | --- | --- | --- |
| Simulation 1 | 3 | 1 | 7 | {1,4,8} | 60 | 150 | 0 |
| Simulation 2 | {3,5,7} | 1 | 7 | 8 | 60 | 150 | 0 |
| Simulation 3 | 3 | {1,2,4} | 7 | 8 | 60 | 150 | 0 |
| Simulation 4 | 3 | 2 | 7 | 8 | {30,60,100,150} | 150 | 0 |
| Simulation 5 | 3 | 2 | 7 | 8 | 60 | {30,60,100,150} | 0 |
| **Simulation 6** | 3 | 2 | 7 | 8 | 60 | 150 | {0,18,30,42,60} |

Each simulated dataset included 300 variables and 40 samples grouped according to two treatments (trt1 and trt2) and two batches (batch1 and batch2). The balanced batch × treatment experimental design included 10 samples from two batches respectively in each treatment group, while the unbalanced design had 4 and 16 samples from batch1 and batch2 respectively in trt1, 16 and 4 samples from batch1 and batch2 in trt2 (see Table 1).

We first generated two base components ***t***^(*trt*)^ and ***t***^(*batch*)^ to represent the underlying treatment and batch variation across samples in the datasets. The samples with trt1 or trt2 in the component ***t***^(*trt*)^ were generated from 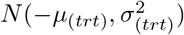 and 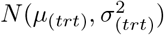 respectively, where 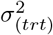 refers to the variability of treatment effect, and similarly for the batch component. We then sampled the corresponding loading vectors ***α***^(*trt*)^ and ***α***^(*batch*)^ from a uniform distribution [–0.3, –0.2] ∪ [0.2,0.3] respectively and scaled them as an unit vector. We subsequently constructed the treatment relevant matrix as ***X***^(*trt*)^ = ***t***^(*trt*)^ (***α***^(*trt*)^)^T^ and similarly for the batch relevant matrix.

We also generated background noise 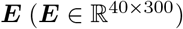, where each element was randomly sampled from *N*(0, 0.2^2^). The final simulated dataset ***X***_*result*_ was constructed based on the treatment, batch relevant matrices and background noise. Starting with ***X***_*result*_ = ***E***, we then added different types of variables, such that:

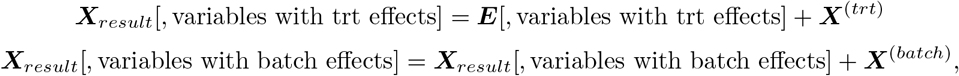

where variables with treatment or batch effects were randomly indexed in the data.

Finally, we simulated a ground-truth dataset that only included the background noise and treatment but no batch effect to evaluate batch corrected datasets.

We simulated different scenarios summarised in Table 2 to verify the influence of different parameters. The scenario indicated in bold is likely to be encountered in real data, where a few variables are relevant to treatment effects, while a large number with batch effects that are stronger and more variable than treatment effects. A subset of variables are influenced by both effects. For the PLSDA-batch analyses, we chose *C* – 1 (or *B* – 1) components associated with treatment (or batch) effects (where *C* or *B* represents the total number of treatment or batch groups) as *C* – 1 (or *B* – 1) components are likely to explain 100% variance in ***Y***, and the number of variables with a true treatment effect (*p*^(*trt*)^) as the optimal number to select on each treatment component in sPLSDA-batch.

#### Case studies

We analysed three 16S rRNA amplicon datasets at the operational taxonomic unit (OTU). Our methods are also suitable for the data considered at any other level of taxonomy. The count data were filtered to alleviate sparsity, then transformed with Centered Log Ratio (CLR) transformation, a pragmatic way to handle both uneven library sizes and compositional structure as (Susin *et al*., 2020). CLR also converts skewed data towards a Gaussian-like distribution.

##### Sponge A. aerophoba

This study investigated the relationship between metabolite concentration and microbial abundance on specific sponge tissues (Sacristén-Soriano *et al*., 2011). The dataset includes the relative abundance of 24 OTUs and 32 samples collected from two tissue types (Ectosome vs. Choanosome) and processed on two separate denaturing gradient gels in electrophoresis. The tissue variation is the effect of interest, while the gel variation is the batch effect.

##### Anaerobic digestion

This study explored the microbial indicators that could improve the efficacy of anaerobic digestion (AD) bioprocess and prevent its failure (Chapleur *et al*., 2016). The microbiota was profiled under various conditions. The dataset includes 231 OTUs and 75 samples treated with two different ranges of phenol concentration (effects of interest). These samples were processed at five different dates, which constituted the batch effect to remove.

##### High fat high sugar diet

This study aimed to investigate the effect of high fat high sugar (HFHS) diet on the mouse microbiome (Susin *et al*., 2020). This dataset includes 419 OTUs and 54 samples treated with two types of diets (HFHS vs. normal) and housed in two different facilities (TRI and PACE). The diet variation is the treatment effect, while the facility variation constitutes the potential batch effect. The actual batch effect in this dataset is weak, and enables to assess whether batch correction methods are able to preserve treatment variation in this context.

The case study datasets are available in GitHub https://github.com/EvaYiwenWang/PLSDAbatch. For the PLSDA-batch analyses, we chose the number of components that explained 100% variance in ***Y*** associated with either treatment or batch effects, and the number of relevant variables to select on each treatment component that yielded the lowest BER from repeated cross-validation with four folds and 50 repeats in sPLSDA-batch.

### Benchmarking and assessment of batch effect removal

We compared our approaches with removeBatchEffect and ComBat that were developed for gene expression data from microarray or RNA-seq and are classical univariate methods to correct for batch effects in the literature. The methods are described in Supporting Information section “Existing methods”.

We next describe several performance measures in removing batch effects while preserving treatment effects between the different methods.

#### Accuracy in simulated data

We assessed the accuracy of identifying variables with a true treatment effect from the batch corrected data using two approaches: 1/ univariate method one-way ANOVA (Law *et al*., 2014) to identify differentially abundant taxa between treatment groups (Benjamini-Hochberg adjusted P-value < 0.05) and 2/ multivariate method sparse PLSDA to select the taxa that discriminate treatment groups. Thereafter, we measured the accuracy of selected variables using Precision 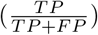, Recall 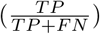 and *F*_1_ score 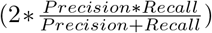, where *TP* is the number of true positives – the variables assigned with treatment effects in the simulation and correctly identified; *FP* the number of false positives - the variables without treatment effects but wrongly identified; FN the number of false negatives - the variables with treatment effects that were not identified. Since in sPLSDA we specified the number of variables to select as the number of variables with a true treatment effect, the Precision, Recall and *F*_1_ score are equal. We thus called this accuracy measure as “multivariate selection” to distinguish from the results from one-way ANOVA (see Table 3).

**Table 3:** Simulation studies: summary of accuracy measures before and after batch correction. The proportion of correctly identified microbial variables with a true treatment effect was assessed with Precision, Recall, F1 score and Multivariate selection score using one-way ANOVA or sPLSDA.

|  |  | Before correction | Ground-truth data | removeBatchEffect | ComBat | PLSDA-batch | sPLSDA-batch |
| --- | --- | --- | --- | --- | --- | --- | --- |
| Balanced | Precision | **0.98** (0.02) | 0.95 (0.03) | 0.94 (0.15) | 0.93 (0.16) | 0.56 (0.25) | 0.86 (0.11) |
|  | Recall | 0.74 (0.10) | **1.00** (0.00) | 0.87 (0.10) | 0.88 (0.10) | **1.00** (0.02) | **1.00** (0.00) |
|  | F1 | 0.84 (0.06) | **0.98** (0.02) | 0.89 (0.12) | 0.89 (0.12) | 0.68 (0.20) | 0.92 (0.07) |
|  | Multivariate selection | 0.89 (0.06) | **1.00** (0.00) | 0.92 (0.07) | 0.92 (0.07) | 0.92 (0.12) | **1.00** (0.01) |
|  |  | Before correction | Ground-truth data | removeBatchEffect | ComBat | wPLSDA-batch | swPLSDA-batch |
| Unbalanced | Precision | 0.52 (0.32) | **0.96** (0.03) | 0.85 (0.18) | 0.80 (0.23) | 0.52 (0.23) | 0.80 (0.14) |
|  | Recall | 0.72 (0.04) | **1.00** (0.00) | 0.86 (0.10) | 0.86 (0.10) | 0.99 (0.03) | **1.00** (0.00) |
|  | F1 | 0.55 (0.21) | **0.98** (0.02) | 0.84 (0.14) | 0.81 (0.18) | 0.65 (0.19) | 0.88 (0.10) |
|  | Multivariate selection | 0.73 (0.05) | **1.00** (0.00) | 0.88 (0.07) | 0.87 (0.08) | 0.89 (0.15) | 0.99 (0.02) |

#### Proportion of explained variance across all the variables

We calculated the proportion of variance explained by treatment, batch effects, and their intersection using the multivariate method partial redundancy analysis (pRDA) in the batch corrected data (Borcard *et al*., 1992, Wang and Lê Cao, 2019).

#### Proportion of explained variance for each variable

The proportion of variance explained by treatment or batch effects for each variable was assessed via *R*^2^ value estimated with one-way ANOVA for each covariate. The *R*^2^ values were then plotted according to treatment or batch on a scatterplot.

#### Alignment scores

To evaluate the degree of mixing samples from different batches in the batch corrected data, we adapted the alignment score originally designed to examine the local neighbourhood of each sample after aligning different groups in single cell RNA-seq data (Butler *et al*., 2018). The alignment score ranges from 0 to 1, representing poor to perfect performance of mixing the samples from different batches after batch effect removal. By applying PCA to the batch corrected data, we calculated a sample dissimilarity matrix with the principal components that explained at least 95% of the total variance. The adapted alignment score is then defined as:

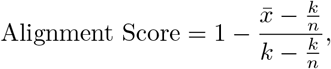

where *k* represents the number of nearest neighbours, and *n* represents the sample size. *x* is the number of each sample’s *k* nearest neighbours that belong to the same batch, and 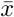 represents the average of all *x*. In our case studies, we chose *k* = 0.1 * *n*, a value that seemed reasonable for the size of our data.

## Results

We benchmarked PLSDA-batch and the two extensions against removeBatchEffect and ComBat, first, on the simulated datasets, then on the three case studies.

### Simulation studies

We measured the accuracy of batch corrected data from different methods applied to the simulated data under different scenarios as shown in the supplements (Figure S1–S6). Here we describe only one scenario that we believe is a representative of real data (*p*^(*trt* & *batch*)^ = 30, simulation 6 in Table 2).

We first considered the proportion of variance explained by treatment and batch effects before and after batch correction across all variables using pRDA. Efficient batch correction methods should generate data with a smaller proportion of batch associated variance and larger proportion of treatment variance compared to the original data. Figure 2**A** shows that there was no intersection shared between treatment and batch variation with a balanced batch × treatment design. All methods successfully removed batch variation, but PLSDA-batch and sPLSDA-batch preserved more proportion of treatment variance than removeBatchEffect and ComBat. In addition, the data corrected by sPLSDA-batch included almost as much proportion of treatment variance as the ground-truth data. With an unbalanced batch × treatment design (Figure 2**B**), we observed that certain amount of variance was shared (intersection) and explained by both batch and treatment effects. Such intersectional variance should exist even in the ground-truth data with no batch effect, as it originates from treatment variation because of the unbalanced design. Unweighted PLSDA-batch and sPLSDA-batch failed in such design, as their corrected data still included a large amount of batch variation (PLSDA-batch) or not included intersectional variance (sPLSDA-batch), while the other methods removed batch variation successfully. The corrected data from removeBatchEffect and ComBat included less proportion of variance explained by treatment but more intersectional variance compared to the ground-truth data. Although wPLSDA-batch corrected data included the largest treatment variance, swPLSDA-batch outperformed all methods with results similar to the ground-truth data.

**Figure 2:**
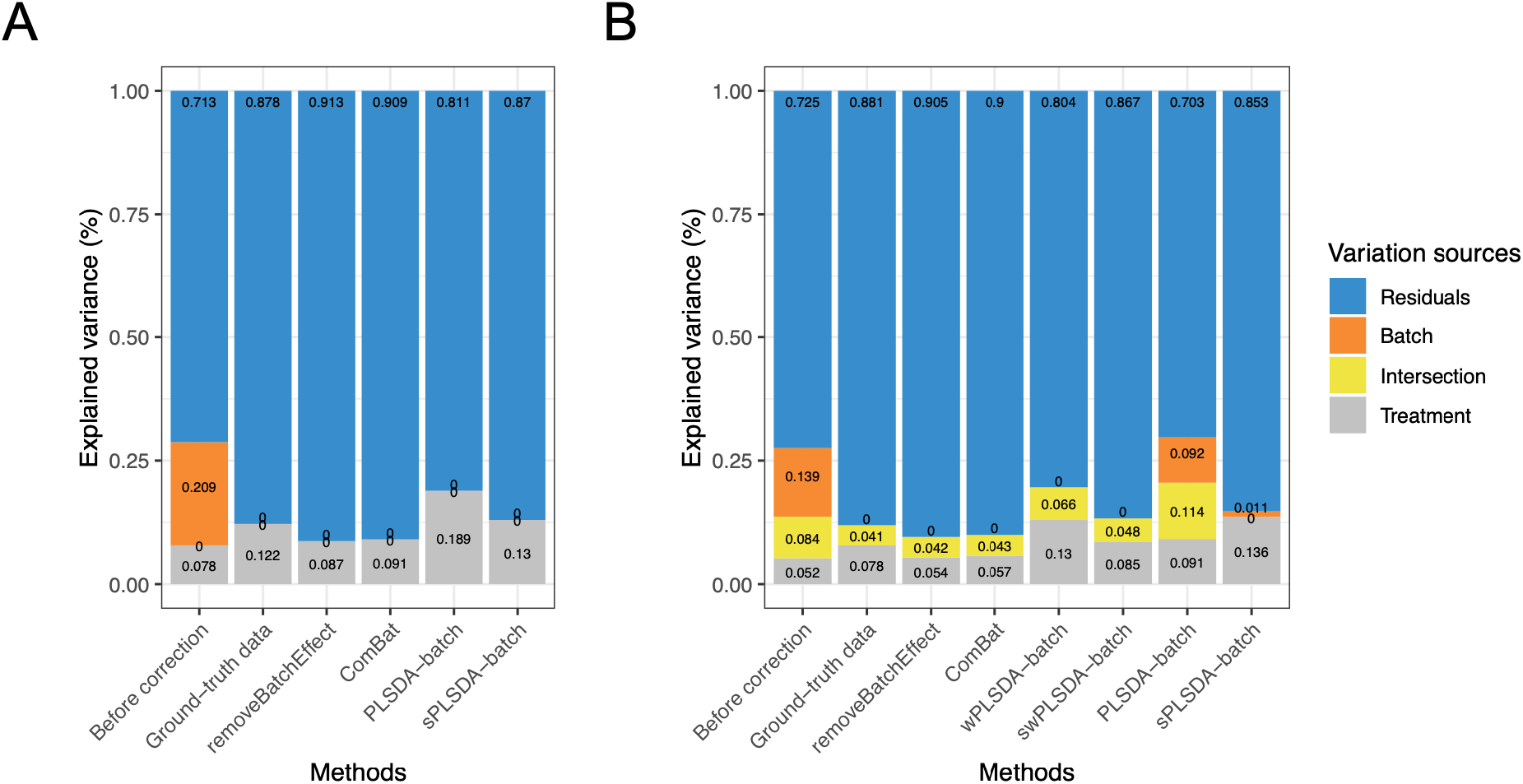
**Simulation studies: comparison of explained variance before and after batch correction** for **(A)** balanced and **(B)** unbalanced batch × treatment designs. The partitioned variance explained by treatment, batch, treatment and batch intersection, and residuals was estimated with pRDA. sPLSDA-batch and swPLSDA-batch performed best in correcting for batch effects as the explained variance was most similar to the ground-truth data that included no batch effect.

We also estimated the proportion of variance explained by treatment and batch effects for each variable respectively using the *R*^2^ value. In the balanced batch × treatment design (Figure 3**A**), the variables assigned with both treatment and batch effects in the corrected data from removeBatchEffect and ComBat presented less proportion of treatment associated variance than in the ground truth data. This result agrees with the pRDA evaluation that these two methods do not preserve enough treatment variation. With PLSDA-batch, variables with only batch effects displayed some amount of treatment variation, but only in the case where the batch effect variability was high (results not shown). sPLSDA-batch outperformed all methods, with results similar to the ground-truth data. In the unbalanced design (Figure 3**B**), variables assigned with both treatment and batch effects were segregated into two groups depending on whether their abundance increased or decreased consistently or not according to the two effects. We observed similar results to those obtained from the balanced design (Figure 3**A**).

**Figure 3:**
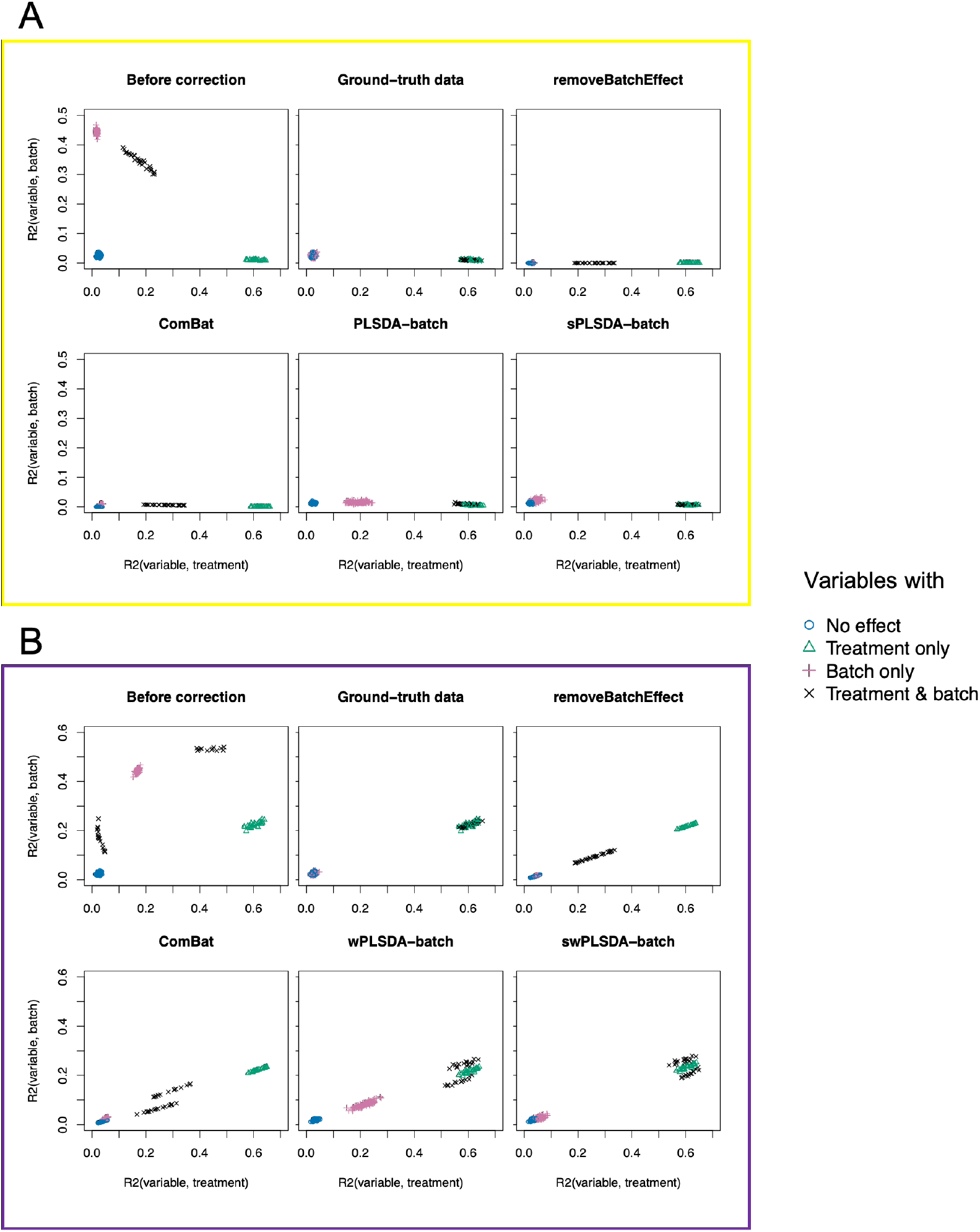
**Simulation studies: R^2^ values for each microbial variable before and after batch correction** for **(A)** balanced and **(B)** unbalanced batch × treatment designs. Each point represents one variable with respect to its fitted R^2^ from a one-way ANOVA with a treatment effect (x-axis) or batch effect (y-axis) as covariate. Colours and shapes indicate the associated effects (batch or/and treatment effects) for each variable. RemoveBatchEffect and ComBat did not preserve enough treatment variation for variables with both treatment and batch effects, while PLSDA-batch and wPLSDA-batch generated spurious treatment variation for variables with batch effect only. sPLSDA-batch and swPLSDA-batch corrected data are the most similar to the ground-truth data that include no batch effects.

When considering the measures of accuracy with univariate one-way ANOVA, we observed that for both balanced and unbalanced designs the corrected data from PLSDA-batch, wPLSDA-batch, sPLSDA-batch and swPLSDA-batch led to higher recall and lower precision than the data from removeBatchEffect and ComBat (Table 3). However, the precision of sPLSDA-batch and swPLSDA-batch was competitive to removeBatchEffect and ComBat for each type of design. Moreover, both versions of weighted and unweighted sPLSDA-batch achieved higher F1 scores and multivariate selection scores than removeBatchEffect and ComBat in each design. The standard deviations of the multivariate selection scores were all smaller than the univariate selection scores for the different corrected data, indicating a better stability of the variables selected by multivariate sPLSDA compared to the one-way ANOVA univariate selection.

We observed similar but higher resolution results of accuracy measures for the other simulation scenarios presented in Figures S1–S6. When the variability of batch effects *σ*_(*batch*)_ increased, the precision of PLSDA-batch decreased dramatically, but the precision of sPLSDA-batch slightly increased and outperformed removeBatchEffect and ComBat in both designs. In all scenarios with a high variability of batch effects (*σ*_(*batch*)_ = 8), PLSDA-batch performed the worst among all the methods. The change of mean (*μ*_(*trt*)_) and variability (*σ*_(*trt*)_) of treatment effects did not largely affect any accuracy measurement. When the number of variables associated either with treatment or batch effects increased, the precision of sPLSDA-batch increased and was slightly higher than removeBatchEffect and ComBat, especially for the unbalanced design. sPLSDA-batch outperformed the other methods in all scenarios except for the case when a large number of variables were influenced by both treatment and batch effects (greater than half the number of variables with treatment effects), resulting in a lower precision but still higher recall than the other two univariate batch correction methods.

### Case studies

#### Numerical performance

We first investigated the variance structure of the batch corrected data using PCA. If batch effects account for the largest proportion of variance in the data, we expect a separation of the samples from different batches on the first component. However, in the sponge data (Figure 4**A**), 21% of the total data variance was explained by the first principal component, which highlighted a strong difference of samples across different tissues (effect of interest). The batch variation accounted for 19% of the total variance in the second component. Thus in this study, batch effects are slightly weaker than the treatment effects.

**Figure 4:**
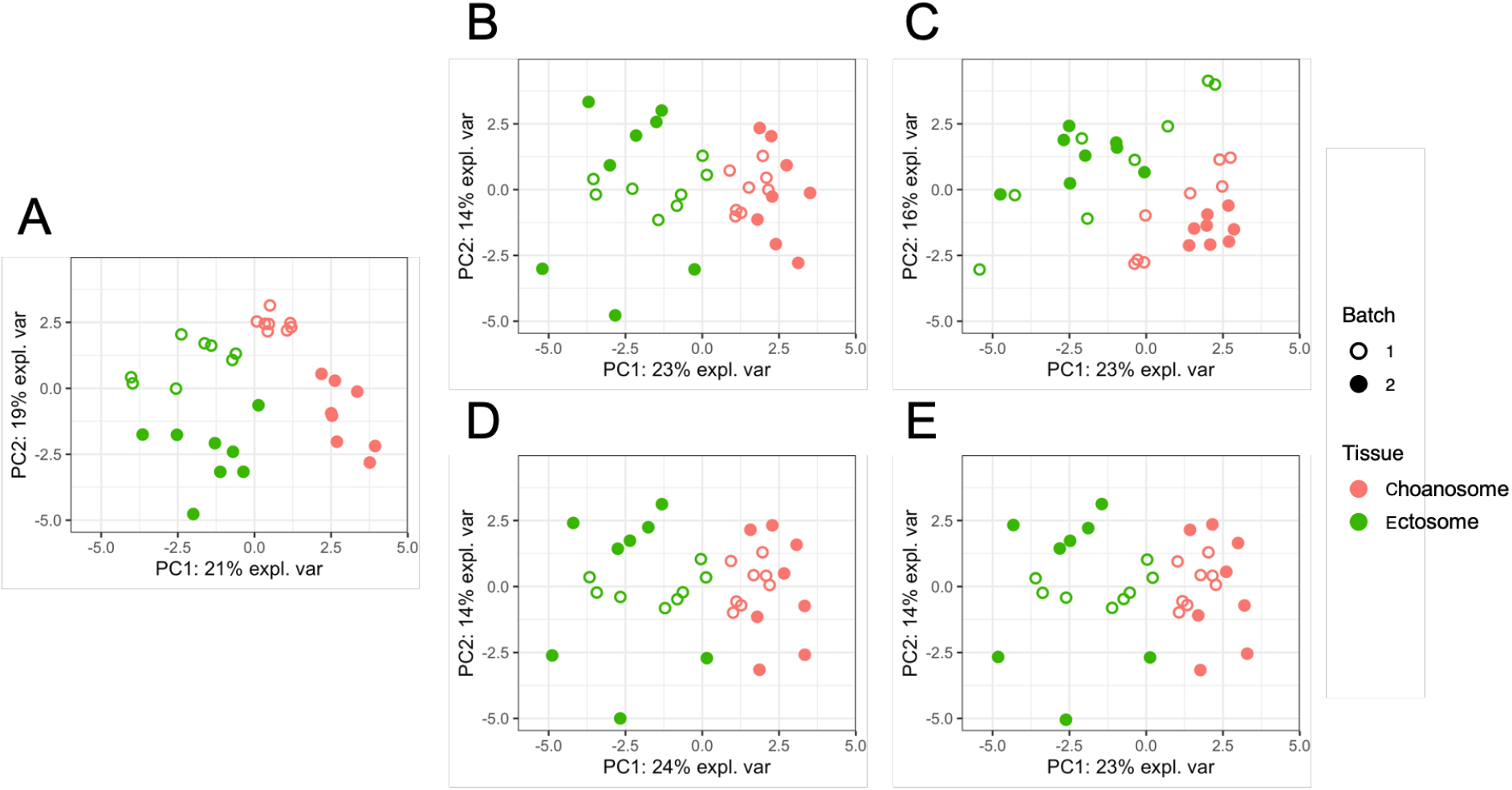
PCA sample plots of the sponge data. **(A)** before or after batch correction using **(B)** removeBatchEffect, **(C)** ComBat, **(D)** PLSDA-batch or **(E)** sPLSDA-batch. Colours represent the effect of interest (tissue types), and shapes the batch types. ComBat did not remove enough batch variation, as samples still present a batch separation within the cluster of Choanosome.

After batch correction, the difference between batches became barely distinct (Figure 4**B-E**), except for ComBat corrected data where a clear separation of the samples from two batches for the Choanosome tissue could still be observed. The variance explained by the first principal component that separated the different tissue types was increased in all of the corrected data, with PLSDA-batch resulting in the highest proportion of variance (24%). We observed similar results in the AD study (Figure S7). When batch variation was not observed on a PCA plot, as for the HFHS data (Figure S8), the proportion of variance explained by the first component (related to treatment effects) before and after batch correction was similar, indicating that treatment variation was preserved. Thus, batch correction methods are still relevant in the case where no batch effect is present.

The alignment scores complement the PCA results especially when batch effect removal is difficult to assess on PCA sample plots. In Figure 5, we observed that the samples across different batches were better mixed after batch correction with different methods than before. In both sponge and AD studies, the data corrected using PLSDA-batch and sPLSDA-batch had higher alignment scores than using removeBatchEffect and ComBat, indicating a better performance in removing batch variation. The ComBat corrected data had the lowest alignment score, which was consistent with PCA that the data still had residual batch variation remaining. In the case of an undetected batch effect, such as HFHS data, the corrected data from PLSDA-batch and sPLSDA-batch had lower alignment scores than those from removeBatchEffect and ComBat.

**Figure 5:**
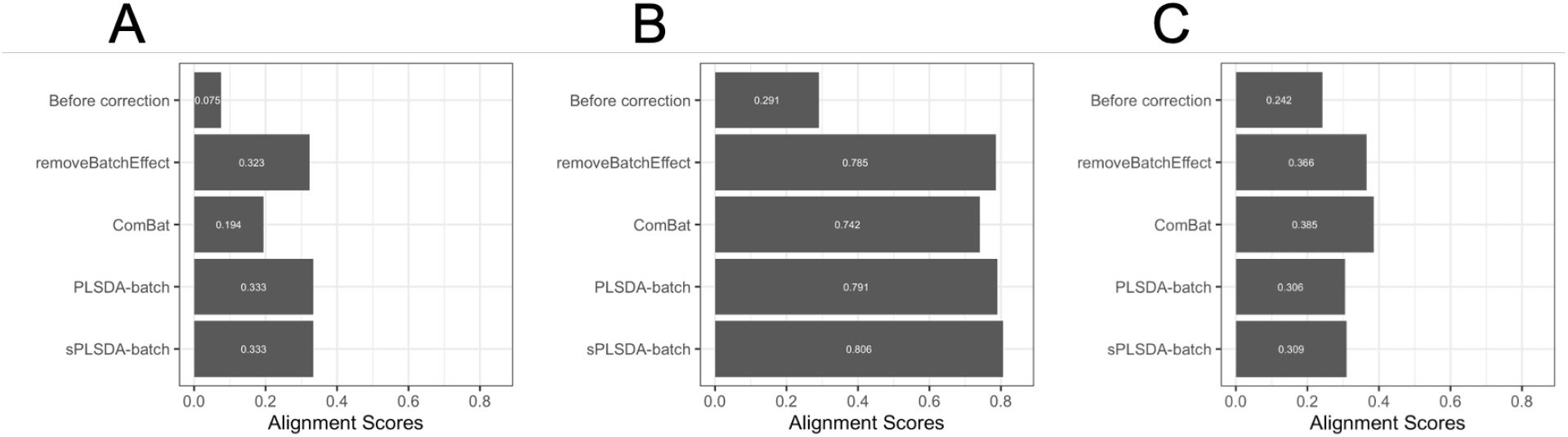
**Comparison of alignment scores** for **(A)** sponge data, **(B)** AD data and **(C)** HFHS data before and after batch correction using different methods. A large alignment score indicates that samples from different batches are well mixed based on the dissimilarity matrix calculated from PCA. For data with strong batch effects (sponge and AD data), our method sPLSDA-batch gave the best performance, while for data with weak batch effects (HFHS data), Combat performed best.

We next focused on estimating the proportion of explained variance by treatment and batch effects globally for the batch corrected data. In the sponge data (Figure 6**A**), the different methods preserved similar proportion of treatment variance and removed all batch variance, with the exception of ComBat that still retained 1.5% of batch variance. In the AD data (Figure 6**B**), we observed a small amount of intersection (0.7%) between batch and treatment associated variance due to the unbalanced batch × treatment design. As the intersection was small, unweighted PLSDA-batch and sPLSDA-batch were still applicable, and thus the weighted version was not used. PLSDA-batch preserved the largest proportion of variance explained by treatment effects, and also the largest proportion of intersectional variance. sPLSDA-batch corrected data led to a slightly higher proportion of treatment variance and an undetectable intersectional variance than the other two univariate methods. In the HFHS data where no batch variation was observed on the PCA plot, we still detected 1.8% of the variance explained by batch effects (Figure 6**C**). The differences of preserved treatment variance and removed batch variance from the different corrected data were small. In addition, the similarity of the results of unweighted PLSDA-batch and sPLSDA-batch in each case study indicated a small batch effect variability.

**Figure 6:**
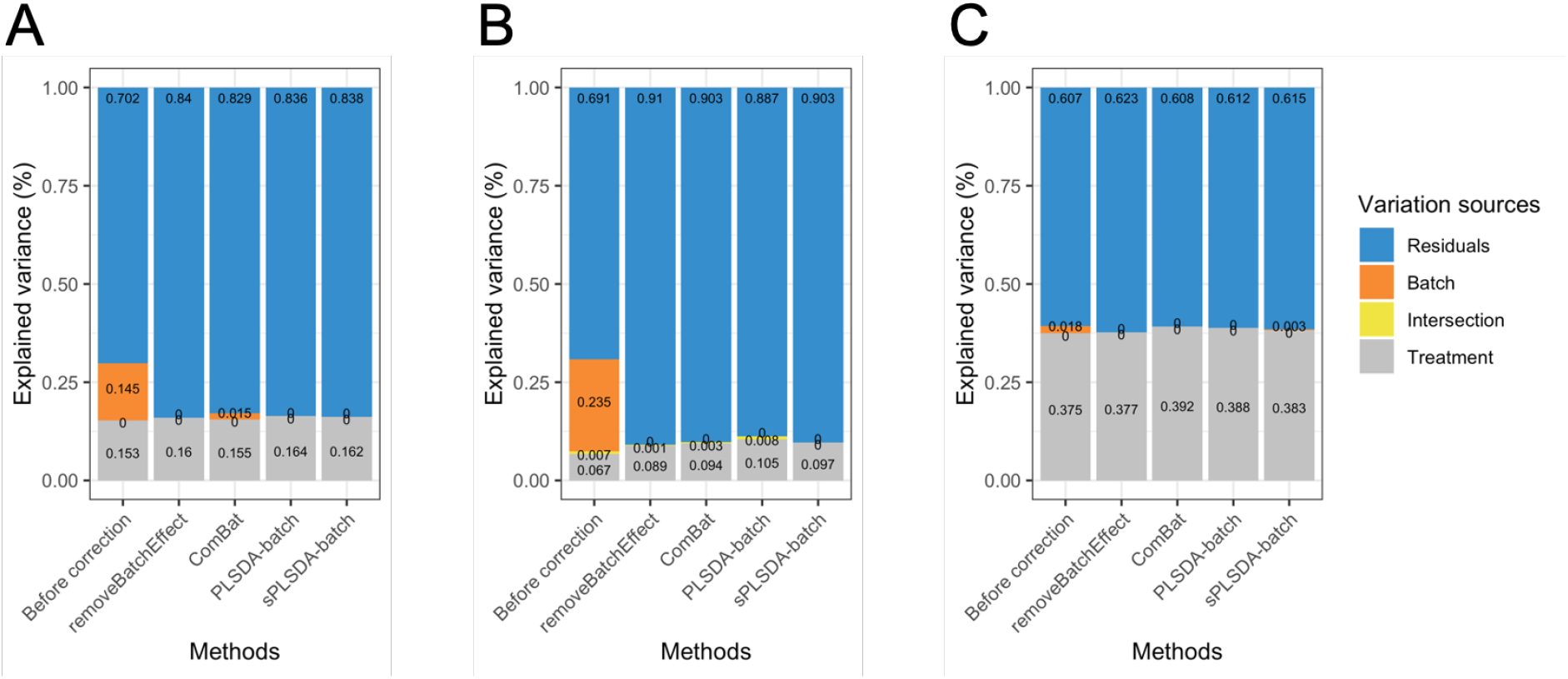
**Explained variance before or after batch correction** for **(A)** sponge data, **(B)** AD data and **(C)** HFHS data. In sponge data **(A)**, ComBat corrected data still included batch associated variance. In AD data **(B)**, sPLSDA-batch corrected data included a higher treatment variance and lower intersectional variance compared to the data corrected from the other methods. In HFHS data with weak batch effects **(C)**, ComBat corrected data preserved the largest amount of treatment variance.

The *R*^2^ values representing the variance explained by batch or treatment effects for each variable estimated with one-way ANOVA are displayed in Figure 7 for the AD study. The corrected data from ComBat still included a few variables with a large proportion of batch variance. When considering the sum of all variables, removeBatchEffect removed slightly more batch variance but preserved less treatment variance than our proposed approaches. The results from the sponge and HFHS data were consistent with the AD data (Figures S9–S10).

**Figure 7:**
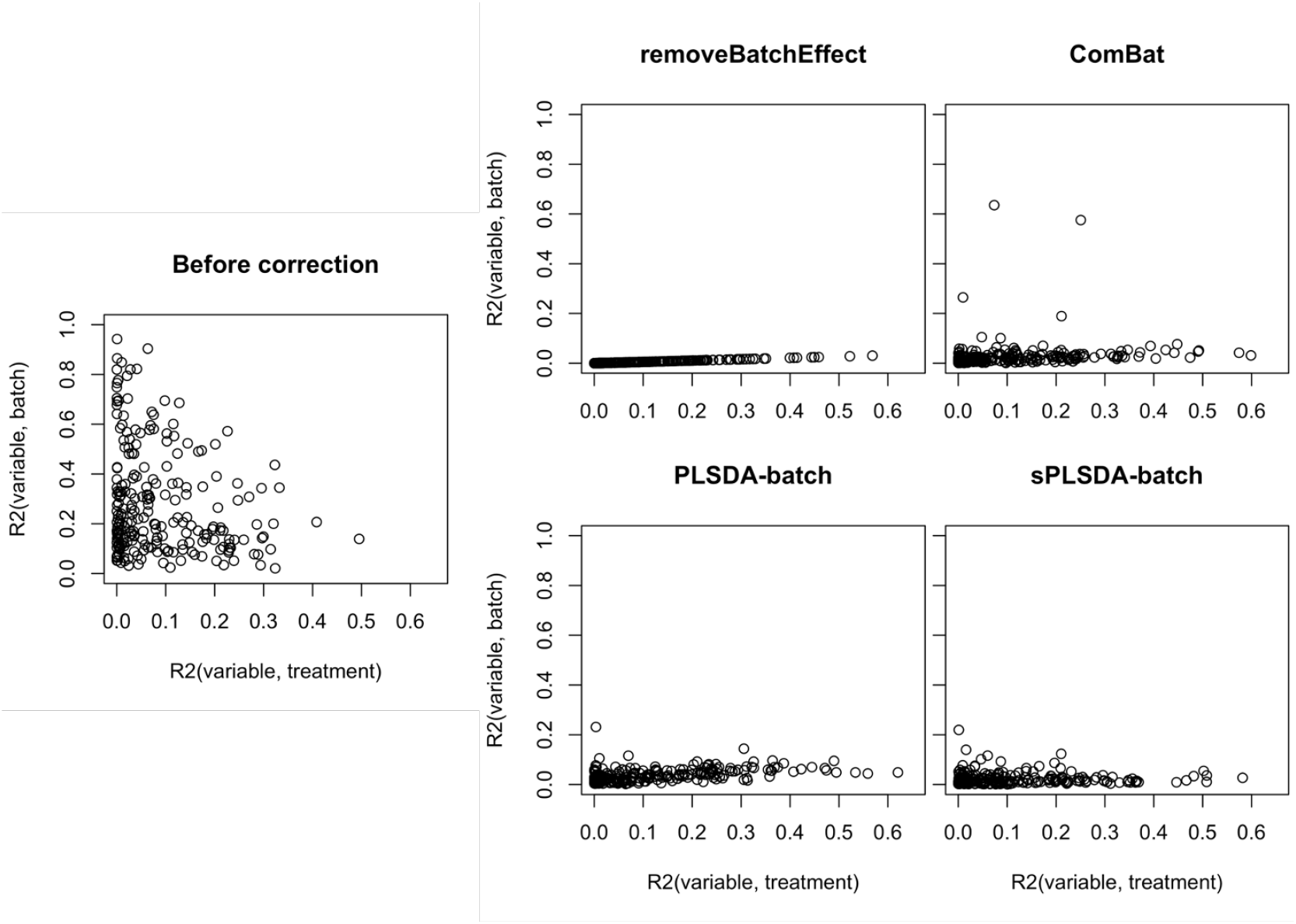
AD study: R^2^ values for each microbial variable before and after batch correction. Each point represents one variable with respect to its fitted *R*^2^ from a one-way ANOVA with a treatment effect (x-axis) or batch effect (y-axis) as covariate. Combat corrected data included some variables with a large proportion of batch variance. Compared to our proposed approaches, removeBatchEffect removed more batch variance.

#### Biological interpretation

We applied sPLSDA to select 20% of the total number of OTUs in the anaerobic digestion and the HFHS studies, but we excluded the sponge study from this analysis since it included a small number of OTUs. We then compared the OTU selections before and after batch effect correction with different methods.

##### Anaerobic digestion

When comparing the variable selections before and after batch correction, two OTUs were uniquely selected in the original uncorrected data, and belonged to the family Methanobacteriaceae and an unknown family of order Clostridiales. Methanobacteriaceae has been reported to be associated with methanogenesis (Granada *et al*., 2018). After batch correction, we observed an overlap of 35 out of the 50 OTUs that were selected from the corrected data with different methods, showing a good agreement among all methods. We also identified 16 OTUs that were only selected from the batch corrected data compared to the original uncorrected data. Among these OTUs, one from the family Porphyromonadaceae was only selected with removeBatchEffect, while two from the family Rikenellaceae and Spirochaetaceae were selected with both removeBatchEffect and ComBat. Two out of these three taxa were from the order Bacteroidales. These taxa have been found to be involved in the degradation of the accumulated volatile fatty acids, propionate production and hydrogenotrophic methanogens (Poirier *et al*., 2016, 2018, Di Gioia *et al*., 2020). Another six OTUs among these 16 were only selected with PLSDA-batch or/and sPLSDA-batch, all of which were from the order Clostridiales. Members of this order have been recognised to hydrolyse a variety of polysaccharides by different mechanisms (Poirier *et al*., 2018). The families of these taxa included Ruminococcaceae (2), Syntrophomonadaceae (1), Peptococcaceae (1) and and two unknown families. All known families have been found to play a key role in AD process, ranging from the degradation of cellulose to acetogenesis, and to syntrophic acetate oxidation (Tian *et al*., 2014, Poirier *et al*., 2016, Wirth *et al*., 2019). To summarise, from the data corrected with our PLSDA-batch and sPLSDA-batch approaches, we identified more taxa within the order Clostridiales, while with removeBatchEffect and ComBat we identified more taxa from the order Bacteroidales. Our approaches selected a larger number of unique OTUs compared to removeBatchEffect and ComBat, and these OTUs are highly relevant to the AD process. This study also shows that our approaches were successful at preserving treatment variation for data that included a strong batch effect.

##### High fat high sugar diet

From the original uncorrected data, one OTU was selected from an unknown family of order *Clostridiales* that was not selected after batch effect correction. When analysing and comparing batch corrected data, 75 out of 80 OTUs were commonly selected among all different methods. We also identified seven OTUs that were uniquely selected by particular methods, including one OTU from the family *Verrucomicrobiaceae* selected from the ComBat corrected data. *Verrucomicrobiaceae* has been reported as a probiotic that can fight the metabolic syndrome (Anhê *et al*., 2016, Shan *et al*., 2019). Another four OTUs were only selected from the data corrected with our PLSDA-batch or/and sPLSDA-batch approaches and belonged to the family *Erysipelotrichaceae, S24-7, Lachnospiraceae* and an unknown family of order *Clostridiales*. All known families have been found to be involved in the regulation of metabolism and immunity (Liu *et al*., 2019, Ma *et al*., 2020), degradation of plant glycan, host glycan, and α-glucan carbohydrates (Zhang *et al*., 2018, Rodríguez-Daza *et al*., 2020) and chronic inflammation of the gut (Zeng *et al*., 2016). To summarise, in the HFHS data that included weak batch effects, over 90% of the selected microbial variables from different batch corrected data were in common with the original uncorrected data. However, our approaches still selected additional OTUs relevant to the HFHS diet compared to removeBatchEffect, ComBat and before batch correction.

## Discussion

Our proposed approach PLSDA-batch aims to estimate and remove batch variation in a multivariate fashion, whilst preserving treatment variation. The batch corrected data can then be used as input in any downstream analyses, such as dimension reduction, visualisation, clustering or differential abundance analysis. The simulation study showed that when the variability of batch effects is high, PLSDA-batch can overfit the component estimation and generate spurious treatment variation. Thus, the sparse version sPLSDA-batch is more suitable in this context to select a subset of microbial variables that are discriminative when estimating treatment components. The weighted variant includes group size weight to handle unbalanced batch × treatment designs and resulted in superior results to the unweighted variants to disentangle correlated batch and treatment effects.

We compared our proposed methods to existing removeBatchEffect and ComBat. These two batch correction methods are univariate and assume each variable has a Gaussian distribution. In addition, ComBat assumes that all variables are affected by batch effects systematically. This assumption does not hold true in practice (Wang and Lê Cao, 2019). Our approach PLSDA-batch has a more relaxed assumption about data distribution compared to removeBatchEffect and ComBat, and thus is more suitable for microbiome data, even after CLR transformation. The multivariate nature of our approach also enables to model the correlation structure between variables and handle non-systematic batch effects.

In the simulation studies, we found sPLSDA-batch and its weighted variant outperformed the other batch correction methods in both balanced and unbalanced batch × treatment designs, when the variability of batch effects was high. Generally, our methods preserved a larger proportion of global treatment variance than removeBatchEffect and ComBat. However, only the sparse variant corrected data with explained variance most similar to the ground-truth data that included no batch effect. Using different types of performance measures to assess the relevance of the OTUs selected, we observed consistent results regarding the ability of sPLSDA-batch to reveal treatment variation (competitive precision and higher recall with one-way ANOVA, and higher multivariate selection score with sPLSDA compared to removeBatchEffect and ComBat). The precision score was also higher than with PLSDA-batch. Similar results were also obtained for the weighted version in the data with an unbalanced design.

In the case studies, PLSDA-batch and sPLSDA-batch performed similarly, however, sPLSDA-batch which includes variable selection selected fewer components than with PLSDA-batch according to the BER criteria. These results confirm that irrelevant variables influence component estimation during the batch effect correction process. Both sponge and anaerobic digestion data included a strong batch effect. For both studies, all performance criteria we used indicated that PLSDA-batch and sPLSDA-batch outperformed ComBat, which removed an insufficient amount of batch variation. The data corrected with removeBatchEffect consisted of similar proportion of batch and treatment variance, but worse alignment scores were obtained compared to our methods. When performing variable selection on the data corrected for batch effects with our approaches, we selected a larger number of unique OTUs relevant to anaerobic digestion than with the other batch correction approaches. Regarding the HFHS data that included a weak batch effect, the batch corrected data indicated a lower alignment score with our methods compared to removeBatchEffect and ComBat. However, the other assessment measures we used suggested that our methods removed sufficient batch variation (Figures 6**C** and S10). Therefore, it is possible that PLSDA-batch and sPLSDA-batch removed more sampling noise, leading to a decrease in total variance of the corrected data and more emphasis on batch variance. In the case of weak batch effect, the alignment scores may not be fully appropriate. For the HFHS study, we observed a large overlap of OTUs when performing variable selection before and after batch correction by the different methods, but data corrected by our approaches selected additional OTUs highly relevant to HFHS diet, suggesting that batch effect correction is still beneficial when batch effects are weak. Due to the limited resolution of taxonomic information with 16s rRNA sequencing, our biological interpretation was limited to family level. Deeper resolution obtained with whole genome sequencing would give more insight into the biological meaning of the additional OTUs that were selected with our approaches.

The framework we present requires pre-defined batch group information. In the case of unknown batch information, such effect can be identified with exploratory approaches such as Principal Component Analysis or clustering methods to assign samples to data-driven batch groups. Despite our proposed weighted version, our methods are still limited in their ability to handle the presence of an interaction effect between batch and treatment on microbial variables, because this interaction is likely to be non-linear. Only methods which account for batch effects, and not correct for them, would be suitable, such as the linear regression (Wang and Lê Cao, 2019). The approaches we propose are linear techniques, where both explanatory and response components are constructed based on a linear combination of variables in their corresponding matrices, and where we model the linear relationship between both components. It is highly possible that variables in microbiome data interact non-linearly, leading to non-linearly dependent components from explanatory and response matrices. Non-linear approaches based on PLS kernel could also be expanded in our framework (Nguyen and Tsoy, 2017).

## Data Availability Statement

An R package ‘PLSDAbatch’ and all analyses are fully reproducible and available at GitHub: https://github.com/EvaYiwenWang/PLSDAbatch.

## Acknowledgments

We thank A/Prof Olivier Chapleur from INRAE for his help in interpreting the variable selection results from the AD data.

## Supporting Information

### Existing methods

**removeBatchEffect** is a location-scale and univariate method. It has been used in a study of human oral microbiome to remove batch effects caused by different experimental times (Wu *et al*., 2016). Let *X_ijcb_* denotes the abundance value for the variable *j* of sample *i* from the treatment group c and batch *b*. removeBatchEffect includes batch effects as covariates and models *X_ijcb_* as:

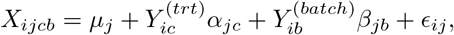

where *μ_j_* is the overall abundance of variable *j*. 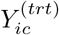 and 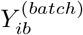 represent the condition of sample *i* in the treatment *c* or batch *b* respectively, and *α_jc_* and *β_jb_* represent the corresponding regression coefficient for the variable *j* in the treatment *c* or batch *b* separately. *ϵ_ij_* is the error term assumed to follow a normal distribution 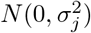. Via removeBatchEffect, we first estimate the batch coefficients and then calculate the batch effect corrected data as 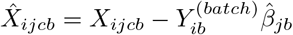.

**ComBat** is a location-scale and univariate method using empirical Bayesian model to estimate parameters. It assumes batch effects are systematic across all variables. ComBat has been applied in a study of human lung microbiome to correct for batch effects caused by different research groups (Hong *et al*., 2018). The abundance value *X_ijcb_* is formulated using the same notations as removeBatchEffect:

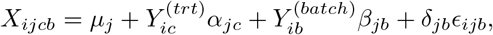

where *δ_jb_* represents the multiplicative batch effect of batch *b* for variable *j*. Both additive (*β_jb_*) and multiplicative batch effects (*δ_jb_*) are modelled in ComBat. The final batch effect corrected data are calculated as 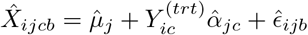.

### Figures

**Figure S1:**
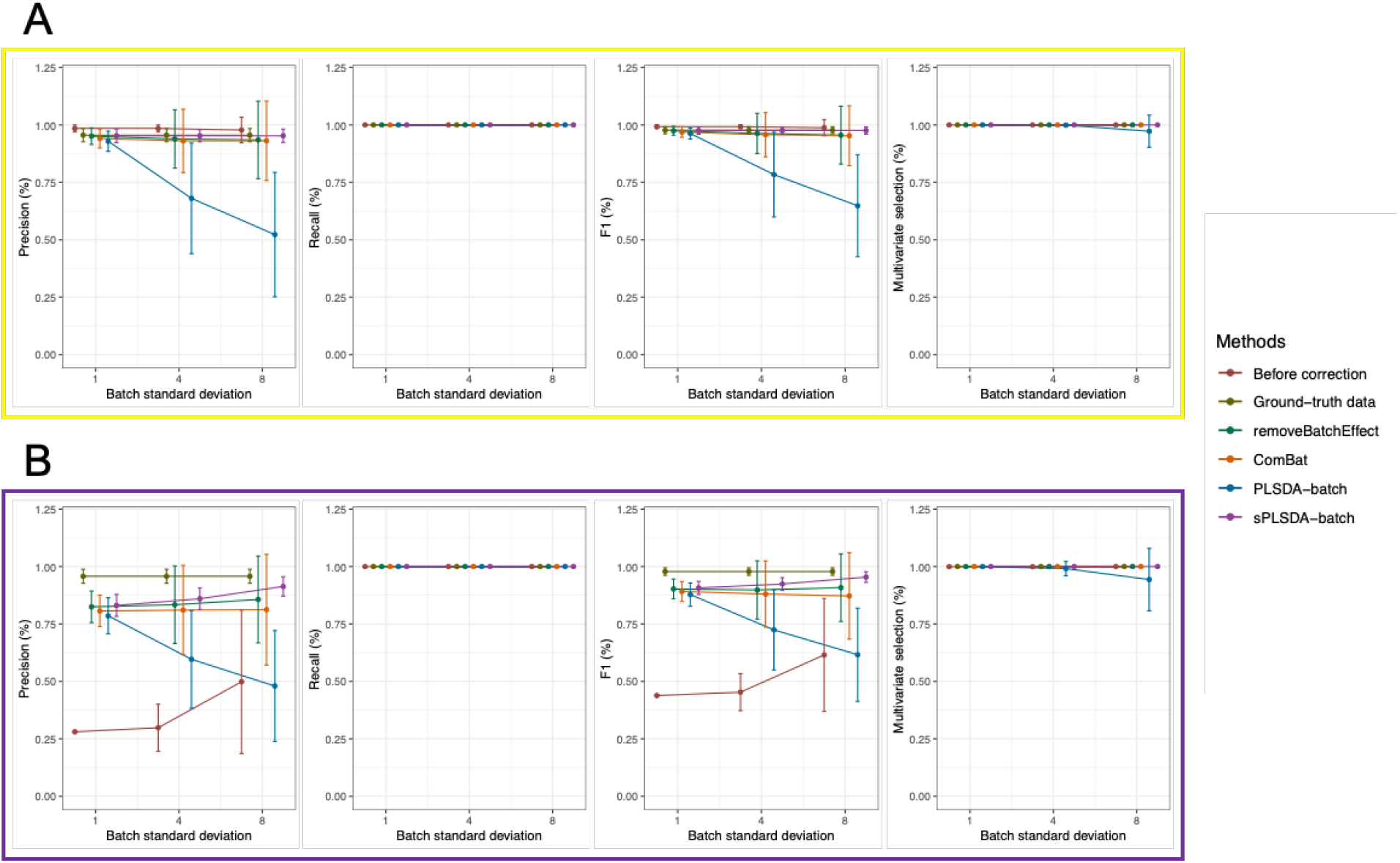
**Simulation 1: summary of accuracy measures before and after batch correction** for the data simulated with different batch effect variability (see Table 2) with **(A)** balanced and **(B)** unbalanced batch × treatment designs. The proportion of correctly identified microbial variables with a true treatment effect was assessed with Precision, Recall, F1 score and Multivariate selection score using one-way ANOVA or sPLSDA. Batch effects were generated with three choices of variability *σ*_(*batch*)_ (x-axis). Each point was averaged over 50 repeatedly simulated data, with error bars indicating estimated sample standard deviations. As *σ*_(*batch*)_ increased, the precision of corrected data from PLSDA-batch dramatically decreased while with sPLSDA-batch slightly increased in both cases of balanced and unbalanced designs. The standard deviation of precision calculated from removeBatchEffect and ComBat corrected data increased with *σ*_(*batch*)_. sPLSDA-batch corrected data slightly outperformed the other corrected data with a higher precision or/and a smaller standard deviation of the precision in both designs. The resulting recall and multivariate selection score were similar among different data. F1 score calculated from the precision and recall therefore displayed the same information as the precision.

**Figure S2:**
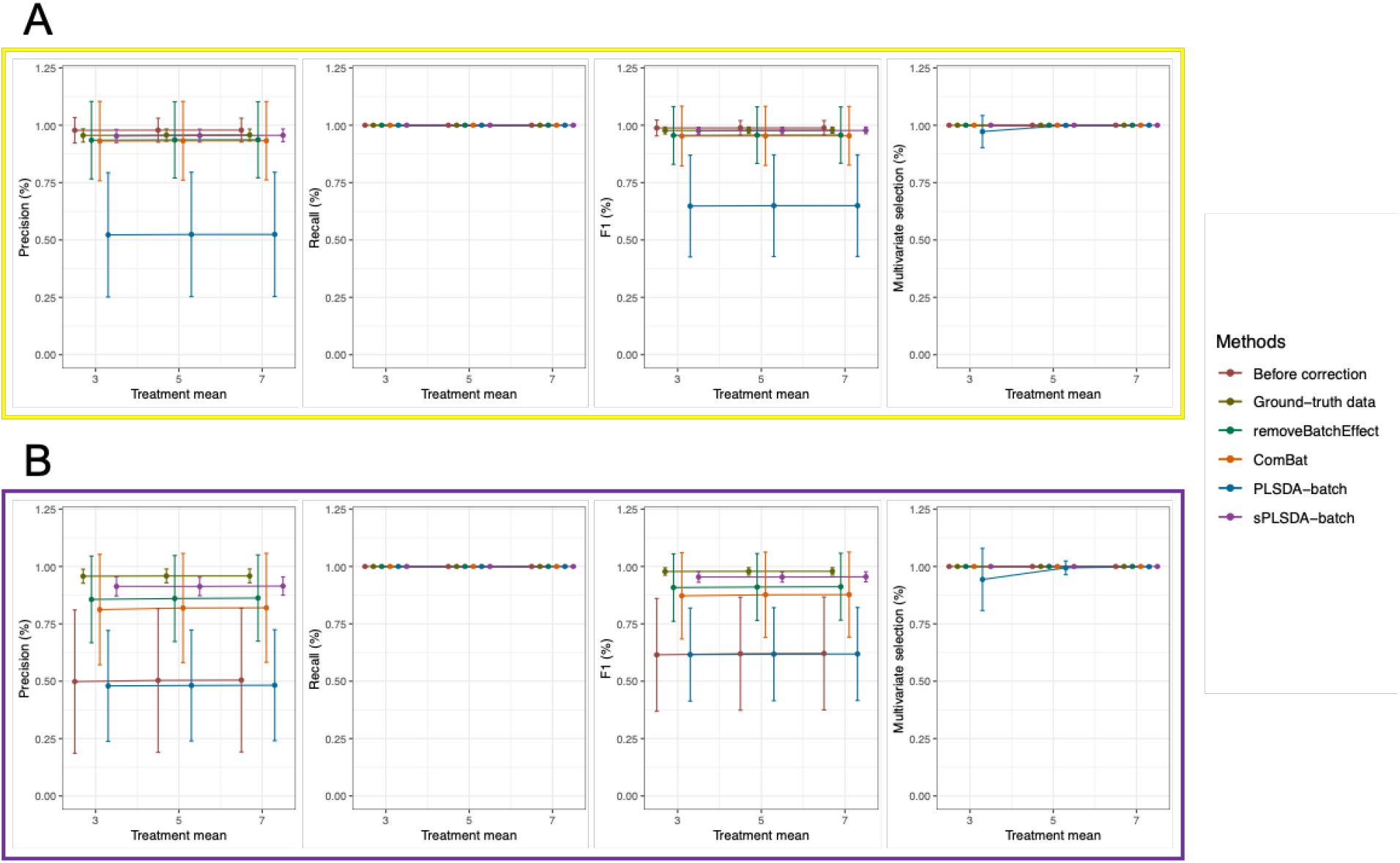
**Simulation 2: summary of accuracy measures before and after batch correction** for the data simulated with different sizes of treatment effects (see Table 2) with **(A)** balanced and **(B)** unbalanced batch × treatment designs. The proportion of correctly identified microbial variables with a true treatment effect was assessed with Precision, Recall, F1 score and Multivariate selection score using one-way ANOVA or sPLSDA. Treatment effects were generated with three choices of sizes *μ*_(*trt*)_ (x-axis). Each point was averaged over 50 repeatedly simulated data, with error bars indicating estimated sample standard deviations. The change of *μ*_(*trt*)_ did not affect the performance of different batch effect correction methods.

**Figure S3:**
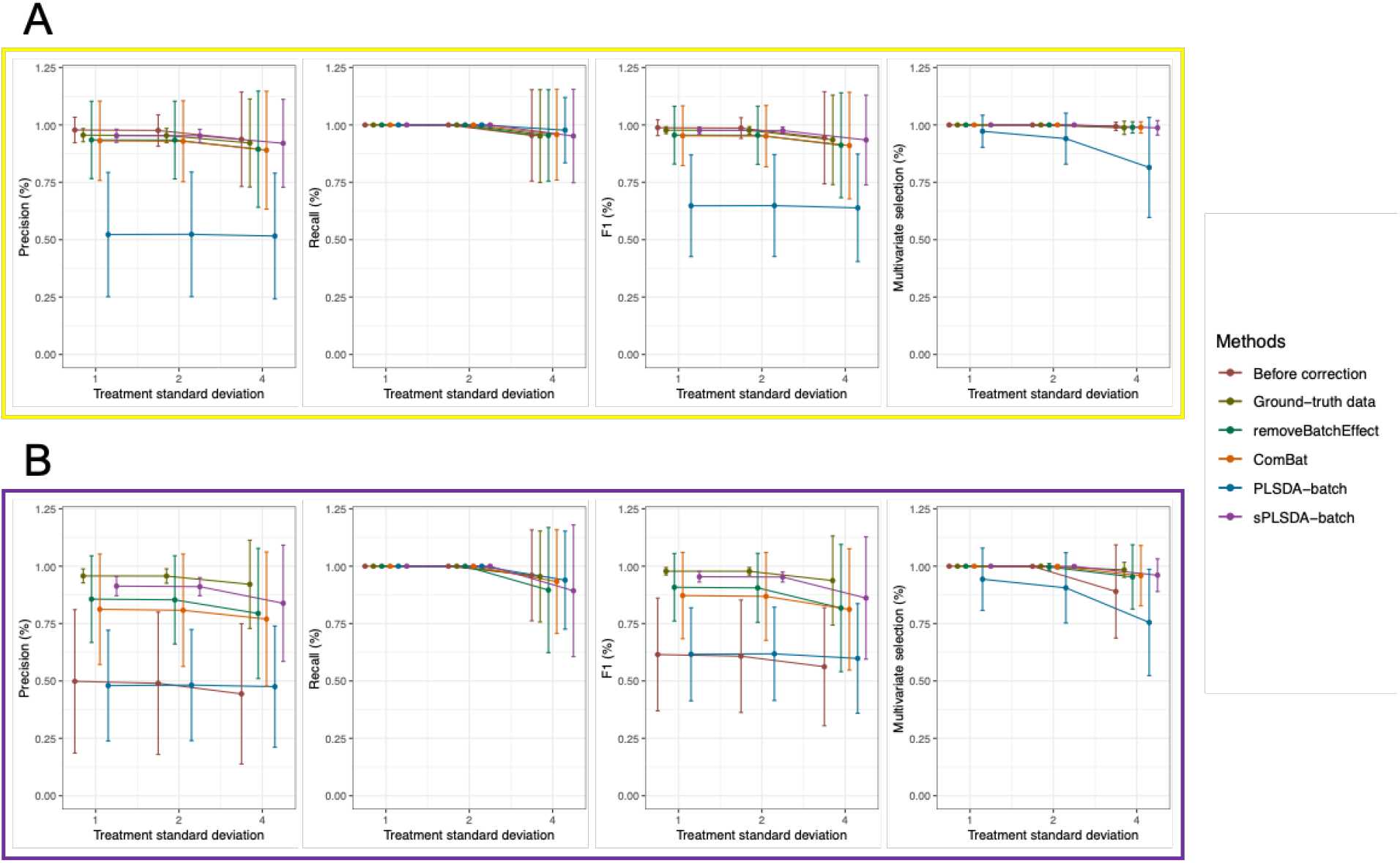
**Simulation 3: summary of accuracy measures before and after batch correction** for the data simulated with different treatment effect variability (see Table 2) with **(A)** balanced and **(B)** unbalanced batch × treatment designs. The proportion of correctly identified microbial variables with a true treatment effect was assessed with Precision, Recall, F1 score and Multivariate selection score using one-way ANOVA or sPLSDA. Treatment effects were generated with three choices of variability *σ*_(*trt*)_ (x-axis). Each point was averaged over 50 repeatedly simulated data, with error bars indicating estimated sample standard deviations. All accuracy measurements of different batch corrected data slightly decreased and their standard deviations increased when the *σ*_(*trt*)_ is larger than 2.

**Figure S4:**
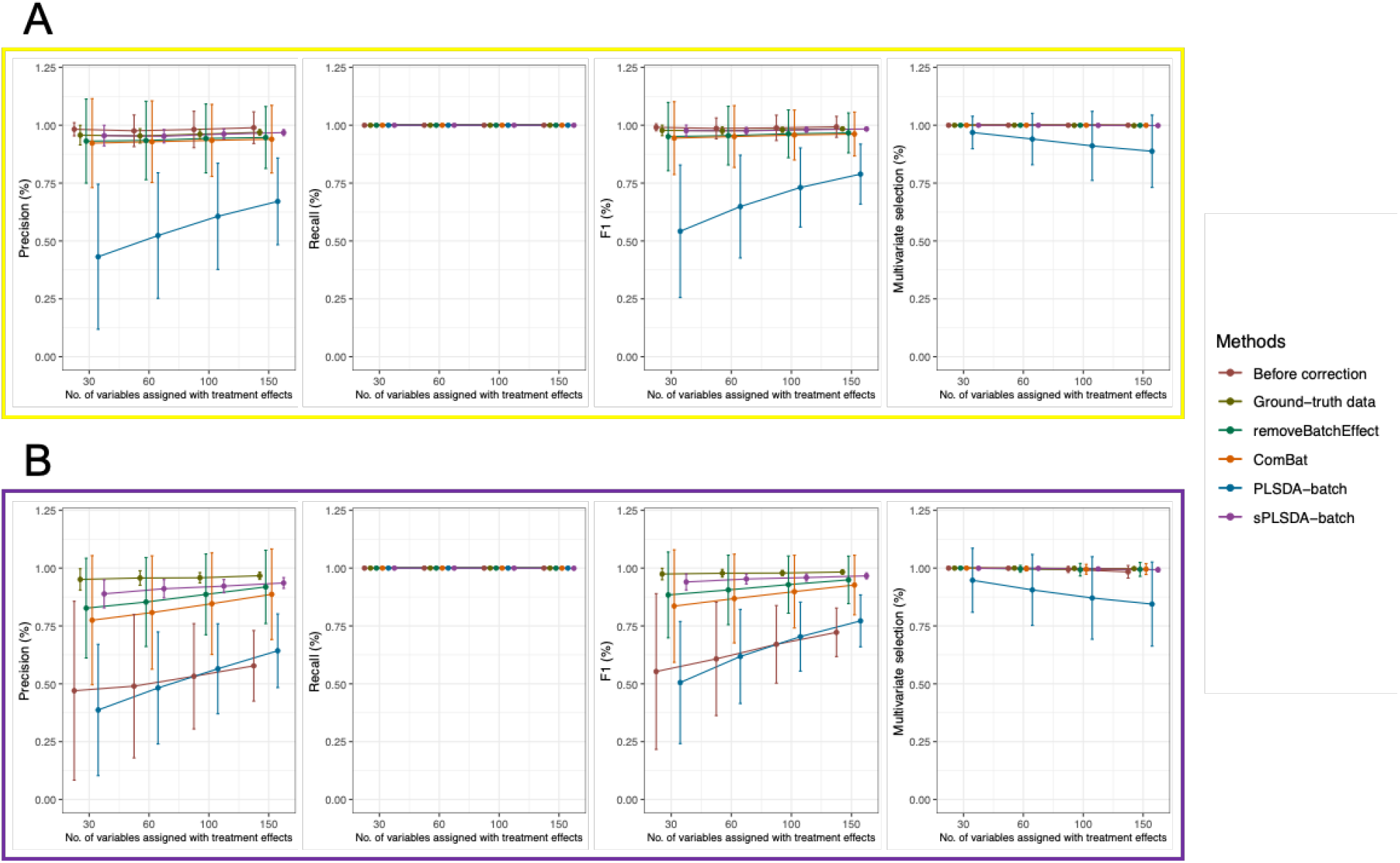
**Simulation 4: summary of accuracy measures before and after batch correction** for the data simulated with different numbers of variables with a true treatment effect (see Table 2) with **(A)** balanced and **(B)** unbalanced batch × treatment designs. The proportion of correctly identified microbial variables with a true treatment effect was assessed with Precision, Recall, F1 score and Multivariate selection score using one-way ANOVA or sPLSDA. Simulated data were generated with four choices of numbers of treatment associated variables *p*^(*trt*)^ (x-axis). Each point was averaged over 50 repeatedly simulated data, with error bars indicating estimated sample standard deviations. The precision of corrected data from different methods slightly increased because of the increase of *p*^(*trt*)^ for the unbalanced design, while similar among different *p*^(*trt*)^ for the balanced deign with an exception of PLSDA-batch corrected data. The multivariate selection scores of different corrected data were similar, except PLSDA-batch corrected data whose multivariate selection score decreased.

**Figure S5:**
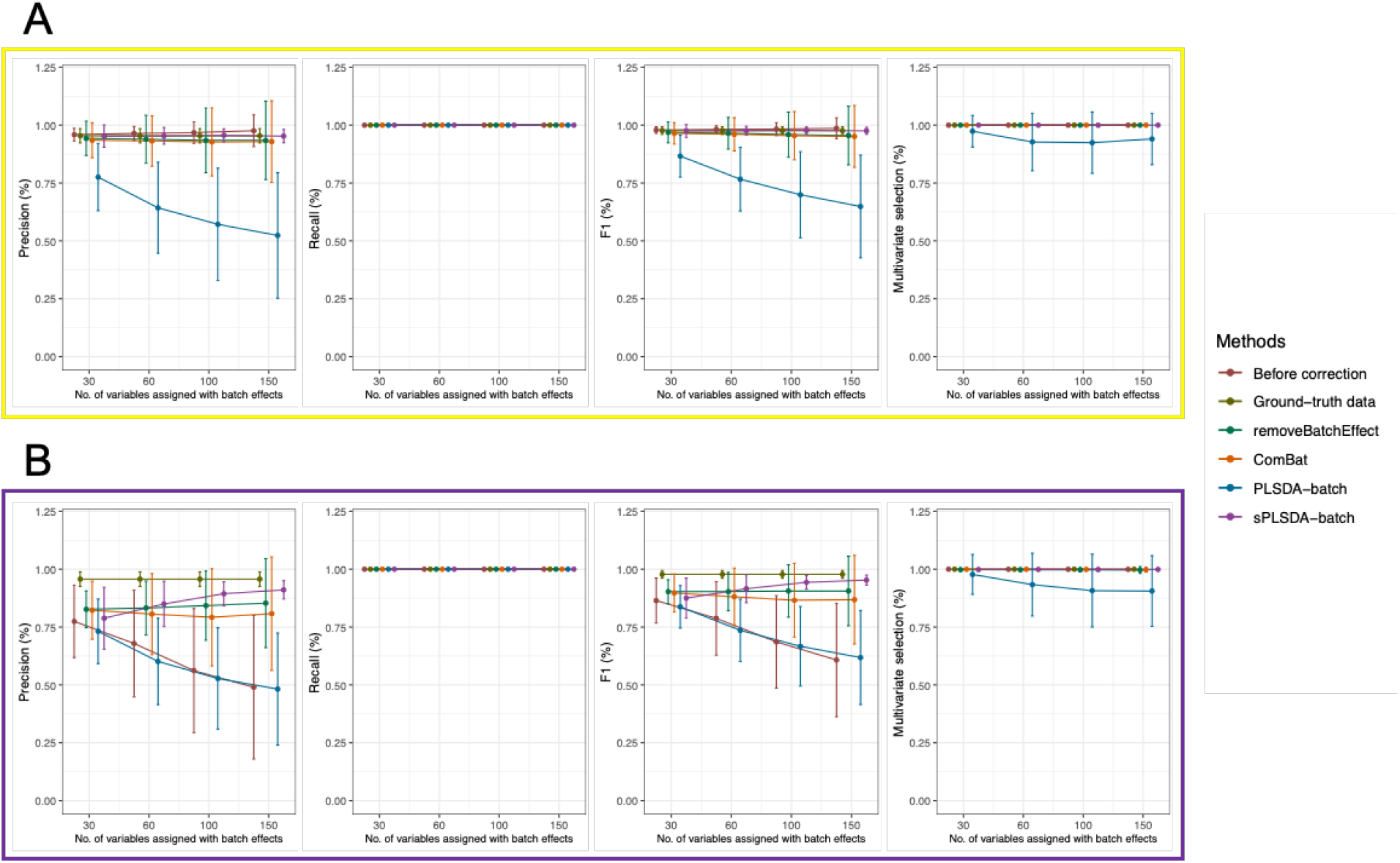
**Simulation 5: summary of accuracy measures before and after batch correction** for the data simulated with different numbers of variables with a true batch effect (see Table 2) with **(A)** balanced and **(B)** unbalanced batch × treatment designs. The proportion of correctly identified microbial variables with a true treatment effect was assessed with Precision, Recall, F1 score and Multivariate selection score using one-way ANOVA or sPLSDA. Simulated data were generated with four choices of numbers of batch associated variables *p*^(*batch*)^ (x-axis). Each point was averaged over 50 repeatedly simulated data, with error bars indicating estimated sample standard deviations. The increase of *p*^(*batch*)^ resulted in an increase of the precision of data corrected with removeBatchEffect, ComBat and sPLSDA-batch, while a decrease with PLSDA-batch for the unbalanced design. The precision of all corrected data and with different *p*^(*batch*)^ were similar for the balanced deign except PLSDA-batch corrected data.

**Figure S6:**
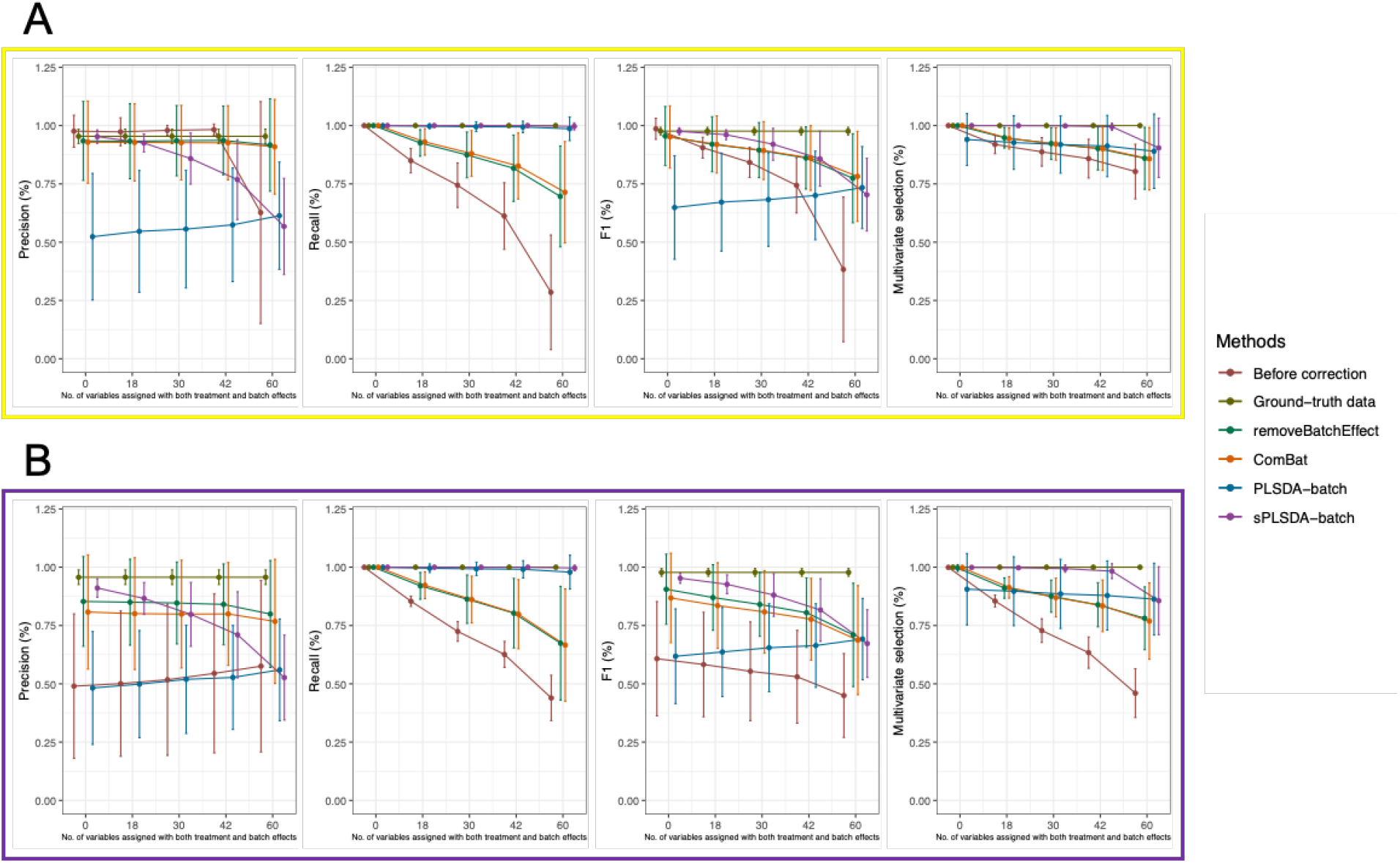
**Simulation 6: summary of accuracy measures before and after batch correction** for the data simulated with different numbers of variables with both treatment and batch effects (see Table 2) with **(A)** balanced and **(B)** unbalanced batch × treatment designs. The proportion of correctly identified microbial variables with a true treatment effect was assessed with Precision, Recall, F1 score and Multivariate selection score using one-way ANOVA or sPLSDA. Simulated data were generated with five choices of numbers of relevant variables with both treatment and batch effects *p*^(*trt* & *batch*)^ (x-axis). Each point was averaged over 50 repeatedly simulated data, with error bars indicating estimated sample standard deviations. When *p*^(*trt* & *batch*)^ was larger than 30 (a half of *p*^(*trt*)^), the precision of data corrected with sPLSDA-batch was lower compared to removeBatchEffect and ComBat, but the recall and multivariate selection score were higher regardless of different *p*^(*trt* & *batch*)^.

**Figure S7:**
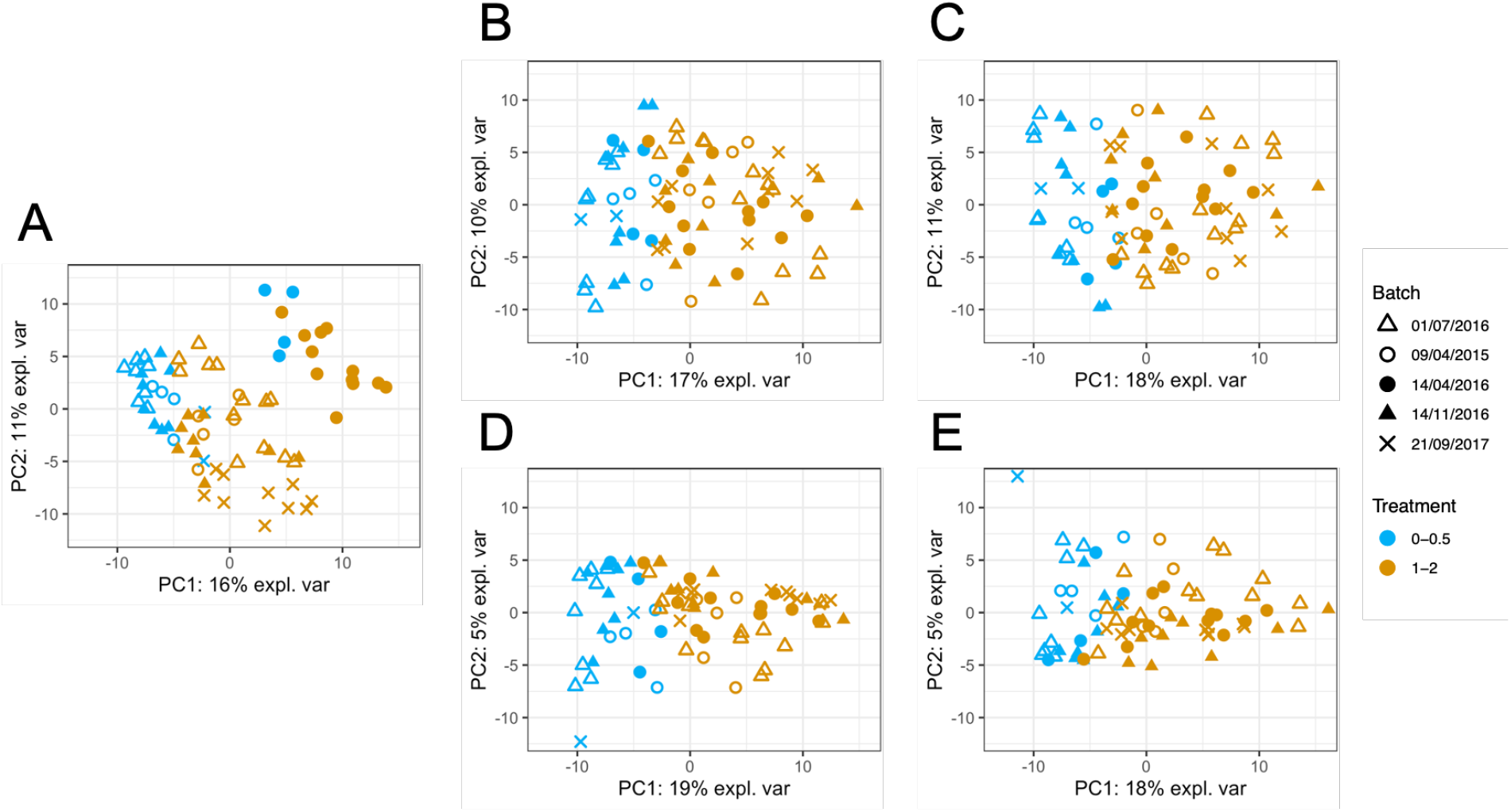
**PCA sample plots of the AD data (A)** before or after batch correction using **(B)** removeBatchEffect, **(C)** ComBat, **(D)** PLSDA-batch and **(E)** sPLSDA-batch. Colours represent the effect of interest (treatment types), and shapes the batch types. The variance explained by the first principal component that separated the different treatment groups was increased in all of the corrected data, with PLSDA-batch resulting in the highest proportion of variance.

**Figure S8:**
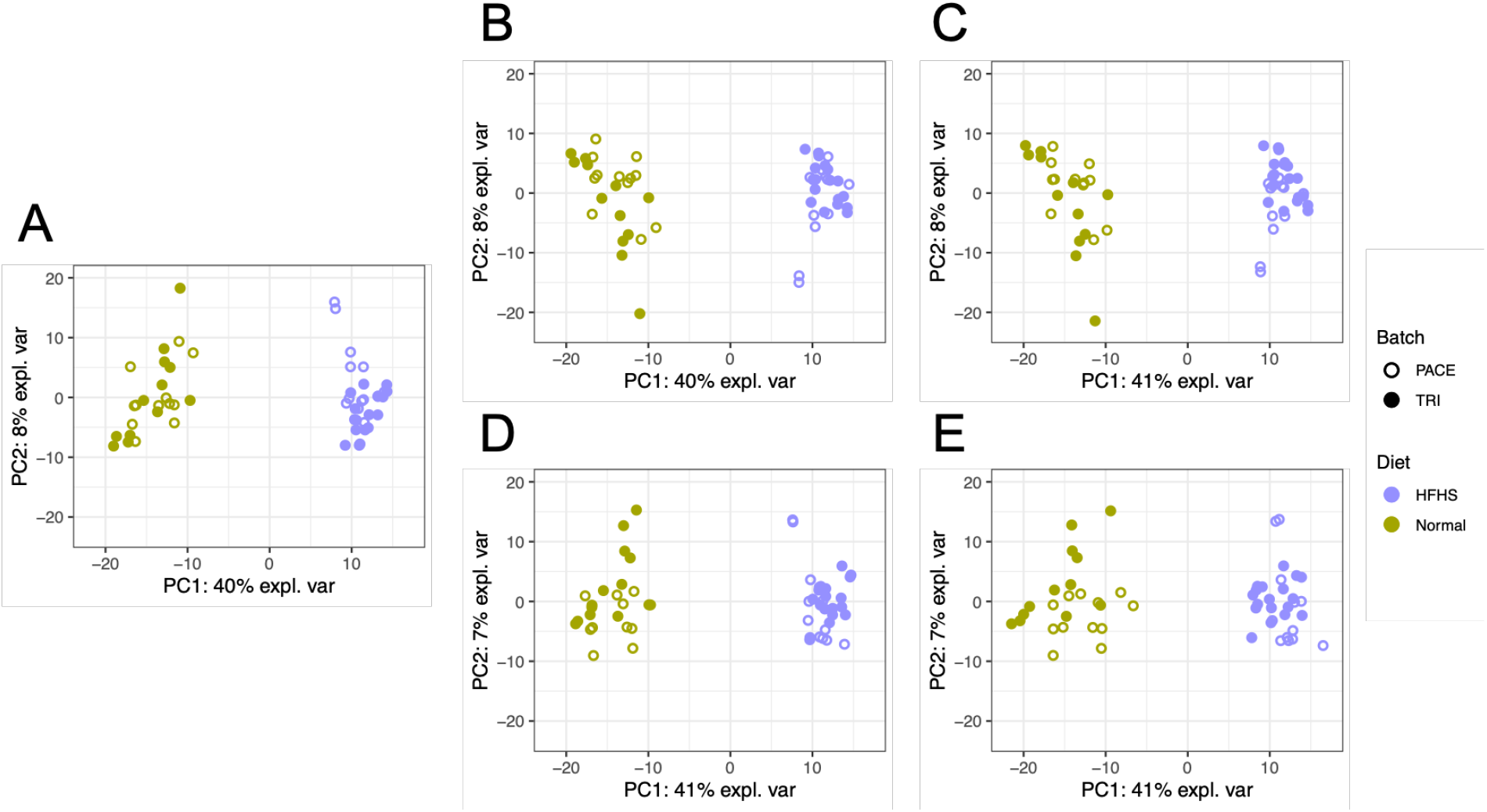
**PCA sample plots of the HFHS data (A)** before or after batch correction using **(B)** removeBatchEffect, **(C)** ComBat, **(D)** PLSDA-batch and **(E)** sPLSDA-batch. Colours represent the effect of interest (diet types), and shapes the batch types. There is no obvious batch variation shown in the data before correction. The proportion of variance explained by the first component (related to diet effects) before batch correction and after was almost the same, indicating a good preservation of treatment variation.

**Figure S9:**
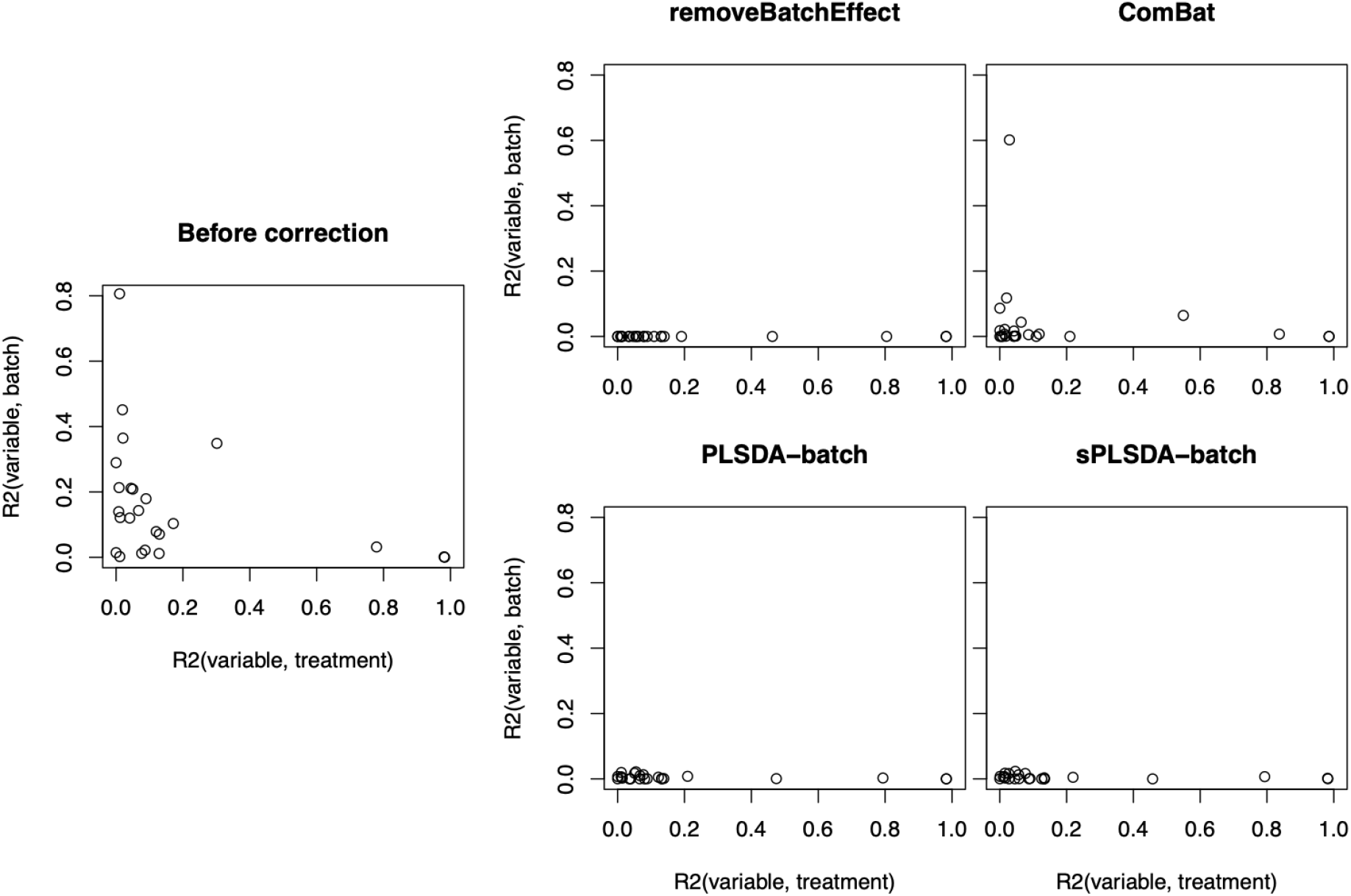
Sponge study: *R*^2^ values for each microbial variable before and after batch correction. Each point represents one variable with respect to its fitted *R*^2^ from a one-way ANOVA with a treatment effect (x-axis) or batch effect (y-axis) as covariate. All methods performed similarly, with an exception of ComBat which included a few variables with batch variance.

**Figure S10:**
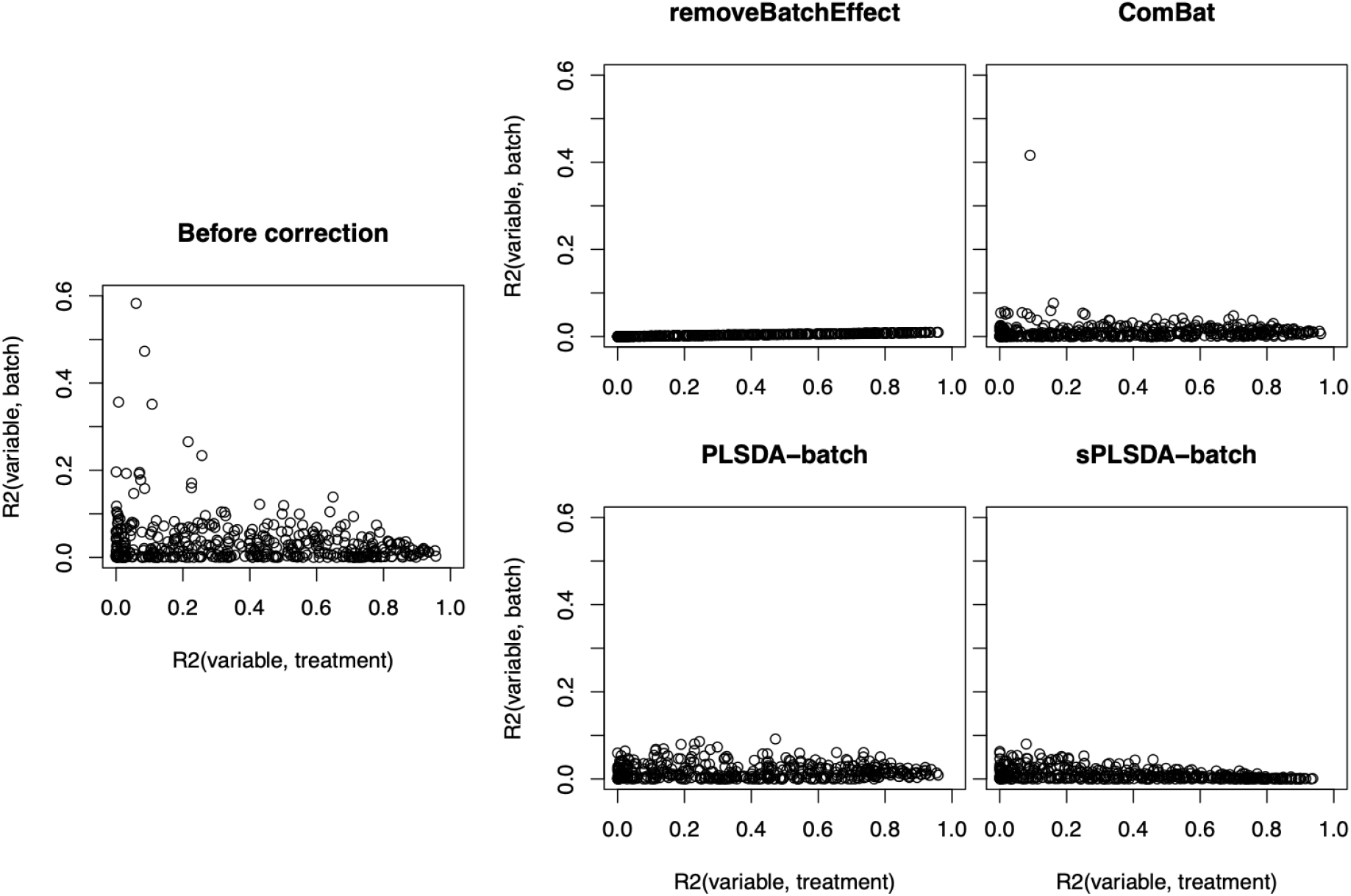
HFHS study: *R*^2^ values for each microbial variable before and after batch correction. Each point represents one variable with respect to its fitted *R*^2^ from a one-way ANOVA with a treatment effect (x-axis) or batch effect (y-axis) as covariate. Combat corrected data included one variable with a large proportion of batch variance. Compared to our proposed approaches, removeBatchEffect removed more batch variance.

